# *Caldicellulosiruptor bescii* regulates pilus expression in response to the polysaccharide, xylan

**DOI:** 10.1101/614800

**Authors:** Asma M.A.M. Khan, Valerie J. Hauk, Mena Ibrahim, Thomas R. Raffel, Sara E. Blumer-Schuette

## Abstract

Biological hydrolysis of cellulose above 70°C involves microorganisms that secrete free enzymes, and deploy separate protein systems to adhere to their substrate. Strongly cellulolytic *Caldicellulosiruptor bescii* is one such extreme thermophile, which deploys modular, multi-functional carbohydrate acting enzymes to deconstruct plant biomass. Additionally, *C. bescii* also encodes for non-catalytic carbohydrate binding proteins, which likely evolved as a mechanism to compete against other heterotrophs in carbon limited biotopes that these bacteria inhabit. Analysis of the *Caldicellulosiruptor* pangenome identified a type IV pilus (T4P) locus encoded upstream of the tāpirins, that is encoded for by all *Caldicellulosiruptor* species. In this study, we sought to determine if the *C. bescii* T4P plays a role in attachment to plant polysaccharides. The major *C. bescii* pilin (CbPilA) was identified by the presence of pilin-like protein domains, paired with transcriptomics and proteomics data. Using immuno-dot blots, we determined that the plant polysaccharide, xylan, induced production of CbPilA 10 to 14-fold higher than glucomannan or xylose. Furthermore, we are able to demonstrate that recombinant CbPilA directly interacts with xylan, and cellulose at elevated temperatures. Localization of CbPilA at the cell surface was confirmed by immunofluorescence microscopy. Lastly, a direct role for CbPilA in cell adhesion was demonstrated using recombinant CbPilA or anti-CbPilA antibodies to reduce *C. bescii* cell adhesion to xylan and crystalline cellulose up to 4.5 and 2-fold, respectively. Based on these observations, we propose that CbPilA and by extension, the T4P, plays a role in *Caldicellulosiruptor* cell attachment to plant biomass.

**IMPORTANCE:** Most microorganisms are capable of attaching to surfaces in part to persist in their environment. Here, we describe that the thermophilic, plant degrading bacterium, *Caldicellulosiruptor bescii*, uses type IV pili to attach to carbohydrates found in plant biomass. This ability is likely key to survival in environments where carbon sources are limiting, allowing *C. bescii* to compete against other plant degrading microorganisms. Interestingly, the carbohydrate that induced the highest expression of pilin protein was xylan, a hemicellulose that is not the majority polysaccharide in plant biomass. Not only do we demonstrate a direct interaction of the pilin with the polysaccharides, but also that cell attachment to polysaccharides can be disrupted by the addition of recombinant pilin and notably by antibody neutralization of the native pilin. This mechanism mirrors those recently described in pathogenic Gram-positive bacteria, and further supports the ancient origins of type IV pilus systems.

## INTRODUCTION

Thermophilic microorganisms capable of hydrolyzing all, or part of lignocellulosic plant biomass have been under considerable interest for biotechnological applications of their enzymes. Of note are cellulolytic microorganisms which produce the enzymes necessary to hydrolyze recalcitrant plant biomass. The Gram-positive, anaerobic, extremely thermophilic genus *Caldicellulosiruptor* employ an array of mechanisms for deconstruction of plant biomass (reviewed in 1). One hallmark of this genus is modular, multifunctional enzymes comprised of both catalytic and binding domains. *Caldicellulosiruptor bescii* is a highly cellulolytic member of the genus (2, 3) capable of attaching to and degrading plant biomass at temperatures as high as 90°C (4). Notably, *C. bescii* is able to degrade insoluble cellulose along with various other plant polysaccharides like xylan (5) and pectin (6), and can grow efficiently on untreated plant biomass with high lignin content (7–9). Modular enzymes with multiple catalytic domains, found primarily in the glucan degradation locus (GDL) (10) diversifies the substrates that these enzymes can hydrolyze (11–15).

In multiple studies, *Caldicellulosiruptor* cells have been observed adhering to plant biomass during growth (4, 9, 16–19), presumably as an adaptation to efficiently degrade lignocellulosic biomass. Given that species from the genus *Caldicellulosiruptor* do not produce a cellulosome, other proteins have been implicated in mediating this attachment. All members of the genus *Caldicellulosiruptor* produce one or more S-layer bound proteins and enzymes (9, 10). Two S-layer located proteins from *Caldicellulosiruptor saccharolyticus* were demonstrated to adhere to cellulose (17). Additionally, S-layer associated enzymes from *Caldicellulosiruptor kronotskyensis* facilitated attachment to xylan when heterologously expressed in *C. bescii* (9). Aside from S-layer located proteins, other potential plant polysaccharide-interacting proteins are also produced by the genus *Caldicellulosiruptor*, including substrate-binding proteins, flagella, a type IV pilus (T4P) and uncharacterized hypothetical proteins which were enriched in a cellulose-bound fraction through a proteomics screen (10). Additional *C. bescii* substrate binding proteins implicated in attachment to plant biomass have also been identified through extracellular protein identification (20), including an expanded proteomics dataset comparing extracellular proteins produced during growth on plant biomass versus crystalline cellulose (21). Both these proteomics studies corroborate the significance of non-catalytic proteins in the process of lignocellulosic plant biomass deconstruction by the genus *Caldicellulosiruptor*.

Structurally unique proteins called tāpirins are another mechanism by which strongly to weakly cellulolytic *Caldicellulosiruptor* species attach to cellulose from plant biomass (19, 22). Genes encoding for tāpirins are located directly downstream of the T4P locus in all cellulolytic *Caldicellulosiruptor* (10, 19, 22), however it remains to be determined if they interact with the T4P.

Protein expression studies using ruminal cellulolytic bacteria identified pilins as cellulose-binding proteins through comparison of binding-deficient mutants versus wild type for *Fibrobacter succinogenes* (23) and *Ruminococcus flavefaciens* (24). Cellulose affinity pull-down assays using extracellular proteins from *R. albus* 8 (25) and extracellular proteome analysis of non-cellulosomal binding deficient *R. albus* 20 mutant also identified pilin-like proteins, further implicating pili in cellular attachment to cellulose (26). Transcriptomic analysis, however, determined that pilin genes encoded by *R. albus* 7 were not differentially expressed on cellulose in comparison to cellobiose, possibly indicating that other polysaccharides act as the inducer for pilin expression (27). Taken together, these studies indicate that T4 pilins from other Gram-positive, cellulolytic bacteria, like the genus *Caldicellulosiruptor*, may facilitate the attachment of cells to cellulose.

Genes required for assembly of a Gram-negative like T4P are fairly widespread throughout the genus *Clostridium* (28, 29), including the pathogens *Clostridium perfringens* (28) and *C. difficile* (30). Major pilins from both *C. perfringens* (31) and *C. difficile* (32) have been demonstrated to play a role in adhesion, and heterologous expression of the *C. perfringens* major pilin gene, *pilA2*, in T4P-deficient *Neisseria gonorrhoea* mutants resulted in attachment to muscle cells (31). Since complemented *N. gonorrhea* were capable of new cell-adherence specificity, but not twitching motility, there appears to be a limitation to the conserved structure and function of pilins. Recently, the ATPases required for twitching motility in *C. perfringens* were found to be upregulated in response to colonization and necrosis of murine muscle tissue (33), supporting the role of *C. perfringens* T4P adherence to muscle cells. Furthermore, *C. difficile* Δ*pilA1* mutants, lacking T4P, were significantly reduced in their ability to attach to epithelial cells (32).

Gram-negative like T4P genetic loci are also present in the genomes of the plant biomass degrading genus, *Caldicellulosiruptor* (10). Among the strongly cellulolytic *Caldicellulosiruptor*, this locus is located upstream of the tāpirins and modular, multi-functional cellulases (10). Available transcriptomics (4, 8, 34) and proteomics data (4, 10, 21, 35) indicate that this T4P locus is strongly upregulated, and that pilin peptides are also present during growth on plant biomass or plant-derived polysaccharides. Considering these data, along with compelling evidence from ruminal and pathogenic Firmicutes that indicate the involvement of T4P in adherence (23-25, 31, 32), we propose that the *C. bescii* T4P plays a role in attachment to plant polysaccharides during plant biomass deconstruction.

## RESULTS

### The <u>Caldicellulosiruptor bescii</u> genome encodes for a Type IV pilus

Based on genome sequence data available for *C. bescii* (4) we confirmed that all essential genes required for assembly of a Gram negative-like type IV pilus (T4P) are present (**Fig. 1A**), including a pilin subunit, pre-pilin peptidase, assembly ATPase, and membrane proteins that recruit the ATPase (29). Similar to T4P locus organization in other Gram-positive bacteria (36) the *C. bescii* T4P locus genes are arranged collinear in an operon, as there are no gaps larger than 71bp between the T4P genes, and only a single hairpin sequence is predicted within the pre-pilin peptidase coding sequence in the T4P locus (**Table S1**). In order to identify putative pilins from the *C. bescii* T4P locus, we screened amino acid sequences for an *N*-terminal pre-pilin cleavage site (GFxxxE) that is post translationally modified by the prepilin peptidase (PilD) prior to pilus assembly (37). We also screened for typical Gram-positive sortase dependent amino acid motifs (LPxTG, 38) however none were identified in the T4P locus. Based on the presence of a pre-pilin cleavage site, we identified five genes (Athe_1872, Athe_1876, Athe_1877, Athe_1880 and Athe_1881) as encoding for putative pilins (**Table 1**). Predicted protein lengths for these range in size from 130aa (Athe_1880) to 277aa (Athe_1872), furthermore, when we analyzed the leader peptide for all five predicted pilins, they all were of variable length, ranging from 5 up to 21 amino acids long (**Table 1**, **Fig. S1**). Based on the total predicted amino acid length of Athe_1880 and Athe_1881, these proteins are typical of T4a pilins (39), however, Athe_1880 has a leader peptide 15 amino acids in length which is not typical for T4a pilins.

**Figure 1.**
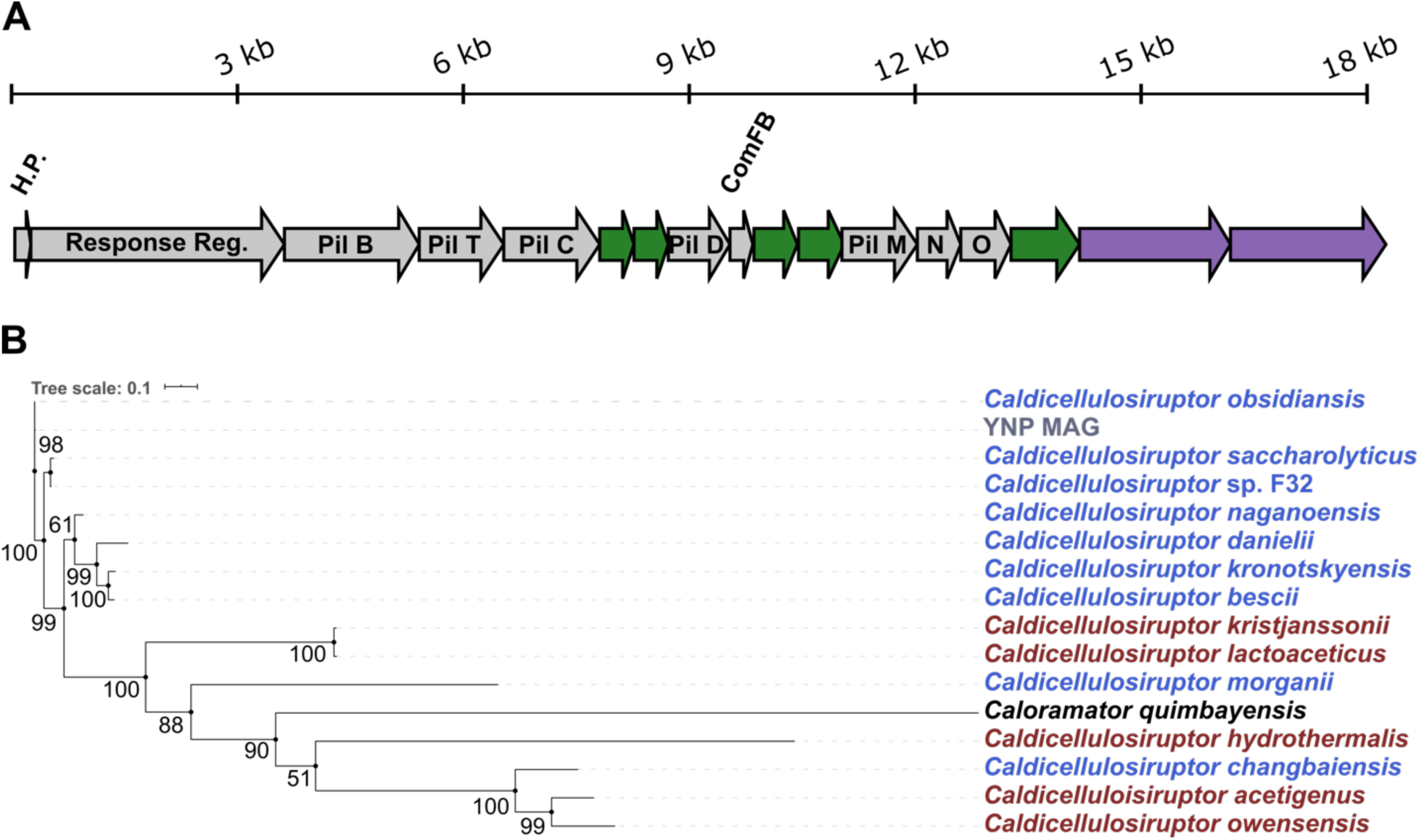
Caldicellulosiruptor Type IV pilus genomic context and evolutionary divergence. (A) *C. bescii* T4P locus organization includes the following genes annotated as (from left to right): H.P., hypothetical protein (Athe_1886); response regulator (Athe_1885); PilB, ATPase (Athe_1884), PilT, ATPase (Athe_1883); PilC, secretion system protein (Athe_1882); PilD, pre pilin peptidase (Athe_1879); ComFB, late competence development protein (Athe_1878); PilM, pilus assembly protein (Athe_1875); PilN, pilus assembly protein (Athe_1874); PilO, pilus assembly protein (Athe_1873). Hypothetical proteins with predicted prepilin-type N-terminal cleavage domains (green arrows): Athe_1881, Athe_1880, Athe_1877, Athe_1876, and Athe_1872 and tāpirins (purple arrows) Athe_1871, Athe_1870. Modified from (10). The scale bar above indicates length in kilo basepairs. (B) Evolutionary analysis of concatenated type IV pilus for the genus Caldicellulosiruptor. A concatenated Maximum Likelihood phylogenetic tree was compiled using IQ-TREE (62). Amino acid sequences homologous to predicted pilins encoded by C. bescii, based on BLASTp were selected. Amino acid sequences were aligned using MUSCLE (60) prior to concatenation. Branches are the consensus of 1000 bootstraps. Dark red indicates strongly cellulolytic, and blue indicates weakly cellulolytic Caldicellulosiruptor species. YNP MAG is a metagenome assembled genome from Yellowstone National Park (41).

**Table 1.** Gene annotations for candidate pilins in *C. bescii*

| Locus tag<br>(Athe_) | IMG<sup>a</sup> | Pfam classification | Pfam ID | Length<br>(aa)<sup>c</sup> | MW<br>(kDa) | LP<br>(aa)<sup>d</sup> |
| --- | --- | --- | --- | --- | --- | --- |
| 1872 | Hypothetical protein | Prokaryotic N-terminal methylation motif | 07963 | 277 | 31.49 | 5 |
| 1876 | Hypothetical protein | NA<sup>b</sup> |  | 175 | 19.91 | 15 |
| 1877 | Hypothetical protein | Prokaryotic N-terminal methylation motif | 07963 | 179 | 19.87 | 5 |
| 1880 | T4P assembly protein PilA | Prokaryotic N-terminal methylation motif | 07963 | 130 | 14.2 | 15 |
| 1881 | T4P assembly protein PilA | Prokaryotic N-terminal methylation motif | 07963 | 133 | 14.99 | 21 |

<sup>a</sup> IMG, Integrated Microbial Genomes database
<sup>b</sup> NA, not assigned
<sup>c</sup> Protein length (aa, amino acids) and MW, molecular weight are for the predicted mature pilin
<sup>d</sup> LP, leader peptide sequence length in apoprotein (see **Fig. S1**)

When comparing the organization of the T4P locus of *C. bescii* to 14 other sequenced *Caldicellulosiruptor* species including a metagenome assembled genome, we observed that their T4P locus organization was highly conserved. Despite this, orthologous pilins shared as little as 35.2 up to 100% percent amino acid sequence identity (**Table S2**), indicating that there may be some evolutionary adaptations in pilins encoded for by strongly cellulolytic versus weakly cellulolytic species (10, 34, 40–42). A phylogenetic tree built from alignment of concatenated pilin proteins from 15 sequenced *Caldicellulosiruptor* species was constructed to assess if there were correlations between cellulolytic ability and the T4P (**Fig. 1B**). Interestingly, the majority of highly cellulolytic species cluster together, however, the pilin proteins encoded by cellulolytic *C. morganii* and *C. changbaiensis* are more divergent (**Table S2**), and cluster with weakly cellulolytic species (**Fig. 1B**). While *C. changbaiensis* was demonstrated to be capable of adhering to cellulose, it is incapable of adhering to xylan (42), indicating that the divergent amino acid sequences may impact adherence to polysaccharides. This difference in the attachment strategies within the *Caldicellulosiruptor* genus lends credence to our primary hypothesis that the T4P, on account of its proximity to modular, multi-domain glycoside hydrolases, plays a role in plant biomass attachment preceding enzymatic deconstruction.

**Table 2.**
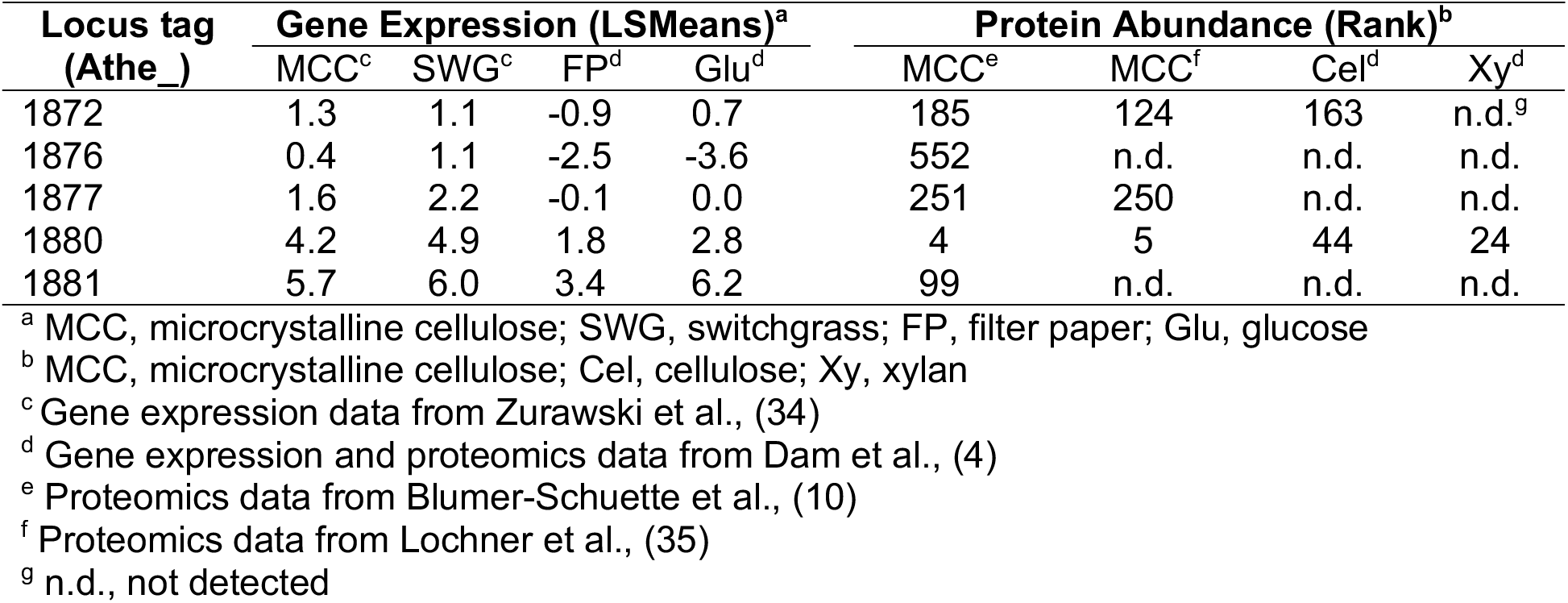
Gene and protein expression data for predicted *C. bescii* pilins

Predicted secondary structures of these putative pilins indicate that these pilins share common regions with the Gram negative T4 pilins which includes an N-terminal alpha helix (α1), an αβ loop separating the α1 helix from the antiparallel β sheets (yellow) in the globular C-terminal domain (**Fig. 2**) (39, 43). Gram negative T4 pilins typically have cysteine residues that define and stabilize the D region (39, 43), however no D region could be identified for *C. bescii* putative pilins as they lack the cysteine residues. Based on the predicted secondary structure and the amino acid length of mature Athe_1880 resembles a T4a pilin, however the major pilin from *Clostridiodes difficile* was also originally predicted to be a T4a pilin on the basis of structural prediction (29), but its solved structure (PDB accessions, 4TSM, 4OGM, and 4PE2) resembled T4b pilins (44).

**Figure 2.**
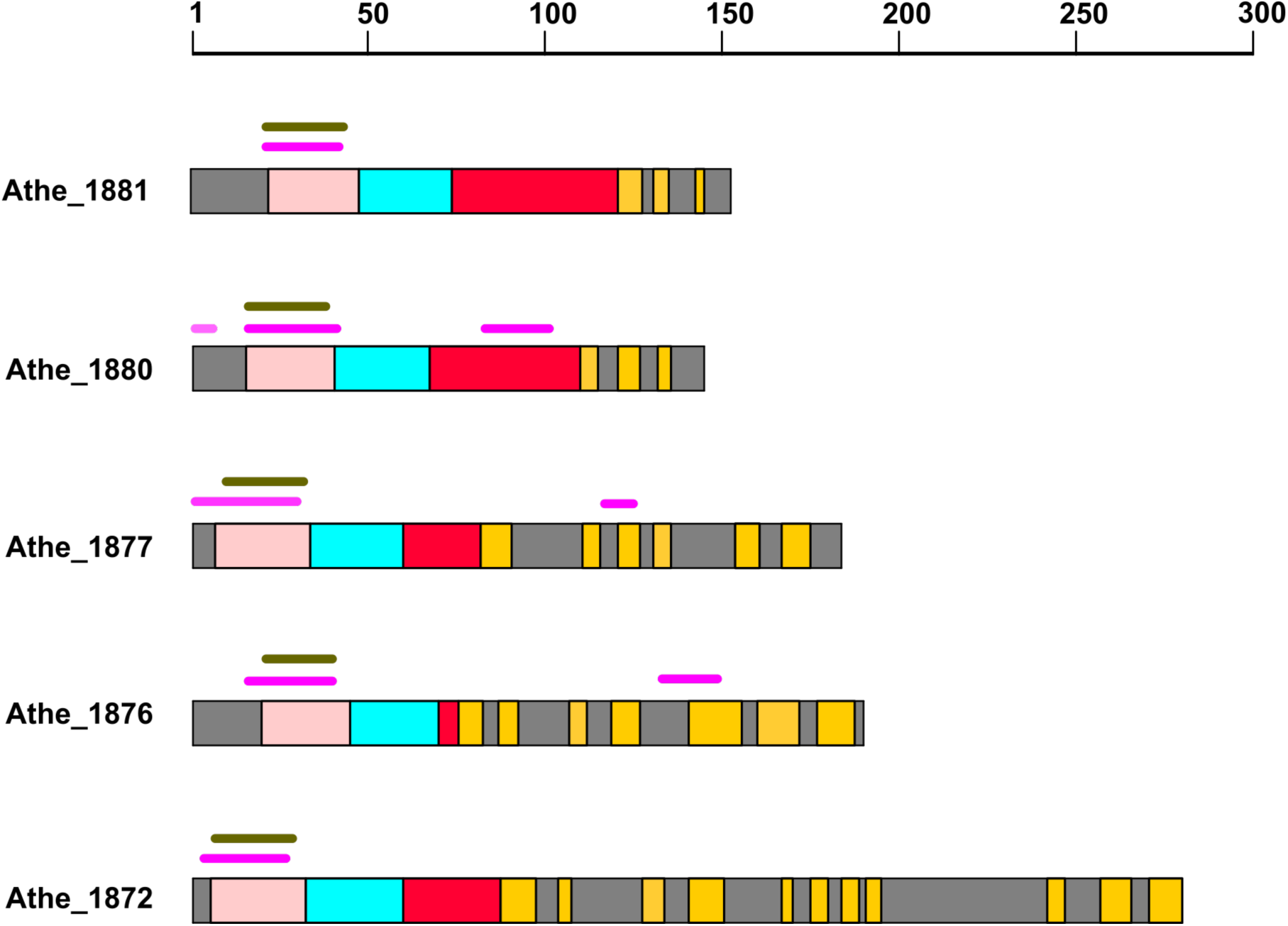
Schematic representation of predicted secondary and tertiary structures for putative *C. bescii* pilins. α1-N helix: peach, α1-C helix: aqua, αβ loop: red, β strands: yellow. Extremely hydrophobic regions are represented by magenta lines above the protein backbone. Transmembrane domains in the α1N region are indicated by olive green lines. Scale bar at the top represents amino acids.

### Athe_1880 is the major pilin (CbPilA) based on transcriptomics and proteomics evidence

We expect that the major *C. bescii* pilin would be highly expressed and would constitute the majority of the T4P structure, therefore we examined publicly available transcriptomics and proteomics data available for *C. bescii*. The major *C. bescii* pilin should either be upregulated as determined by transcriptomics data, or enriched as peptide fragments in proteomics data. Datasets from three independent comparative transcriptomic studies of *C. bescii* grown on switchgrass versus glucose (8), filter paper versus glucose (4) and microcrystalline cellulose versus switchgrass (34) respectively indicated that Athe_1880 and Athe_1881 are the most highly upregulated genes among the candidate pilins (**Table 2**). Proteomics data for protein abundance on microcrystalline cellulose confirmed that Athe_1880 is the most abundant of all of the candidate pilins across three independent sets of proteomics data (4, 10, 35). These data were further corroborated by a recent study on the extracellular proteome of *C. bescii* where Athe_1880 was found to have a fold change greater than 2x on complex substrates like xylan, switchgrass and Avicel compared to simple substrates like xylose, glucose and cellobiose (21). Given both the transcriptomic and proteomic evidence, we propose that Athe_1880 (CbPilA) is the major pilin represented in the T4P filament of *C. bescii*. We then sought to produce soluble, recombinant CbPilA (rCbPilA) using rational design, informed by predicted secondary protein structures (**Fig. 2**) in order to produce truncated rCbPilA lacking the highly hydrophobic α^-1^N region (**Fig. S2**).

### Xylan is the key inducer of CbPilA

Previously published transcriptomics and proteomics data confirmed that *CbpilA* gene expression and protein production were upregulated when *C. bescii* was grown on polysaccharides or plant biomass versus mono- or disaccharides (4, 8, 10, 21, 34, 35). Based on this evidence it is natural to assume that cellulose would be the main polysaccharide regulating the T4P operon. Since the other representative plant polysaccharides had yet to be tested, we sought to examine if hemicellulose polysaccharides played any role in the regulation of the T4P by monitoring the presence of extracellular CbPilA in samples harvested from batch cultures. Polyclonal antibodies were generated against rCbPilA for this purpose, and we confirmed that this antibody was specific for native CbPilA, in comparison to three predicted pilins from *C. bescii* (**Fig. S3**).

In addition to carbohydrate induction of pilin production, we also tracked CbPilA production over time using immuno-dot blots. *C. bescii* cultures were cultured on 5 different plant polysaccharide-related carbohydrates: cellulose, pectin, glucomannan, xylan, and xylose and sampled during early, mid, and late exponential phase. Xylan induced a 10-fold higher amount of CbPilA production (7.7 pg cell^-1^) compared to glucomannan (0.75 pg cell^-1^) or xylose (0.51 pg cell^-1^) at late exponential growth (**Fig. 3**). Furthermore, the amount of CbPilA protein quantified also increased in a growth phase dependent manner for all sugars tested, with no detectable CbPilA present during early exponential phase for growth on glucomannan and xylose. The most dramatic increase was noted during growth on xylan, with over a 3.5-fold increase in CbPilA from early to late exponential growth (**Fig. 3**). Interestingly, while cellulose is the major component of plant biomass, it was the least effective polysaccharide for inducing T4P production during growth on a defined medium, and extracellular CbPilA was below measurable limits, as was also the case for pectin (data not shown). Given that xylan induced the highest levels of CbPilA protein, we concluded that it is in fact, xylan, rather than cellulose that is the main inducer of CbPilA.

**Figure 3.**
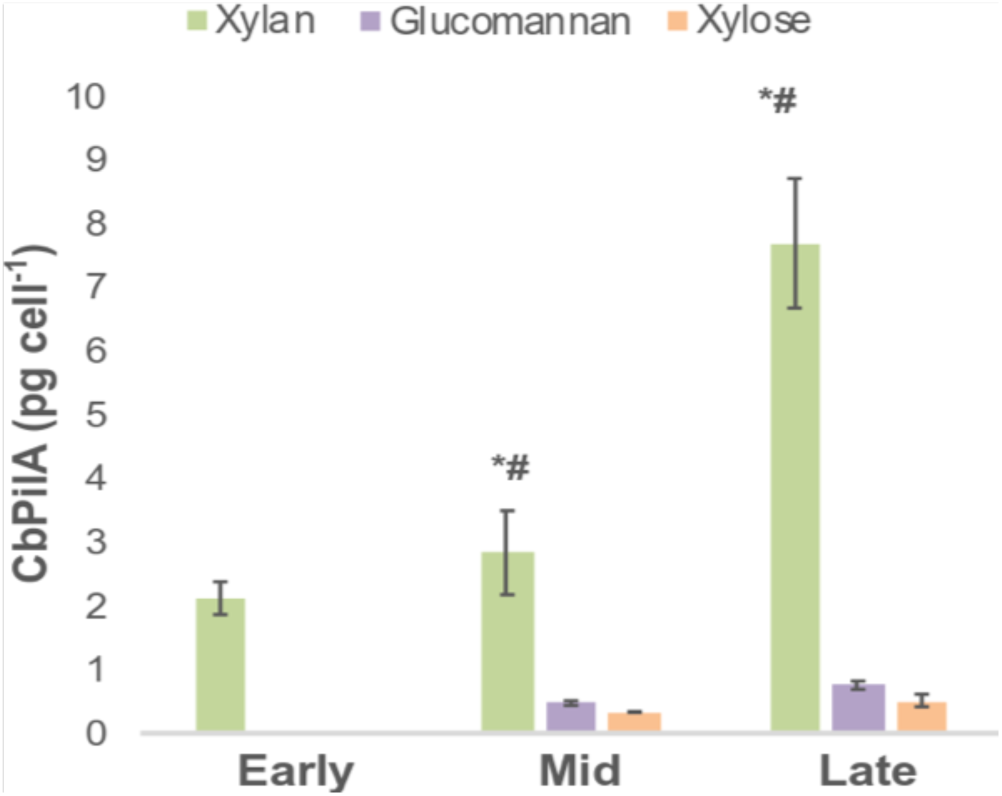
Detection of CbPilA protein levels during growth on representative plant polysaccharides. Normalized CbPilA concentrations are plotted against the y-axis during early, mid and late exponential phases of growth. CbPilA concentrations were determined relative to that of the rCbPilA standard. Standard curves (R squared= 0.98) were built using dilutions of purified rCbPilA (1 ng µl^-1^, 10 ng µl^-1^, 50 ng µl^-1^, 100 ng µl^-1^ and 200 ng µl^-1^) in Image Lab 5.2.1 (Bio-Rad). Each bar is the average from three independent replicates (± standard error), normalized to the number of cells blotted for each time point. * p-value < 0.05 when compared to mean CbPilA production during growth on glucomannan, # p-value < 0.05 when compared to mean CbPilA production during growth on xylose. Representative plant polysaccharides include xylan (green), glucomannan (purple) and xylose (orange). No CbPilA was detected during growth on glucomannan and xylose during early exponential phase.

### rCbPilA interacts with polysaccharides at elevated temperatures

Given that xylan was the main inducing polysaccharide for CbPilA production, we tested for an interaction between rCbPilA and insoluble carbohydrates, using polysaccharide pulldown assays. It is common practice to use incubation temperatures lower than physiological temperature when testing thermostable proteins (45, 46), however no interaction was noted when rCbPilA was incubated with xylan or microcrystalline cellulose at 25°C (**Fig. 4**). When the incubation temperature was increased to a physiologically relevant temperature (75°C), we observed a direct interaction of rCbPilA with insoluble xylan as well as microcrystalline cellulose (**Fig. 4**). This observation is likely the result of increased flexibility of rCbPilA, potentially exposing polysaccharide binding sites. An absence of an rCbPilA protein band in the negative control supports that this interaction is specific for the tested polysaccharides (**Fig. 4**). To validate the specificity of this assay, we tested a recombinant *C. bescii* tāpirin (**Fig. S4A**). As expected, the tāpirin protein was capable of adsorbing to microcrystalline cellulose at 25°C, as was previously observed (19, 22), however a denser band was observed after incubation at 75°C (**Fig. S4B**), similar to our observations with rCbPilA. Densitometry analysis also confirmed a specific interaction of the tāpirin with microcrystalline cellulose (**Fig. S4B**) whereas the density of the protein band in the xylan bound fraction was no different than that of the negative control (**Fig. S4C**).

**Figure 4.**
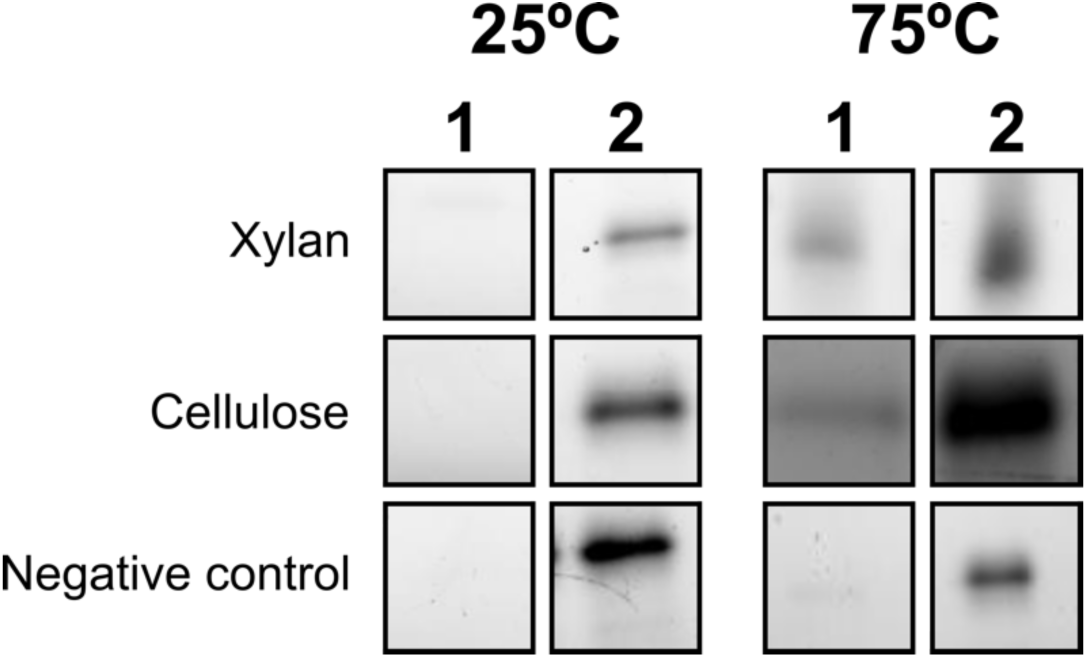
Polysaccharide pulldown assay using rCbPilA. Recombinant CbPilA was incubated with insoluble microcrystalline cellulose or xylan at room temperature (25°C) or physiological temperature (75°C), after which bound and unbound protein fractions were separated for SDS-PAGE. 1, bound protein fraction; 2, unbound protein fraction. Negative control includes rCbPilA and buffer, only. SDS-PAGE images are representative of the results from assays run in triplicate.

### CbPilA is localized to the *C. bescii* cell surface

Although immuno-dot blots confirmed the production of CbPilA protein by *C. bescii* cells (**Fig. 3**), we wanted to confirm that CbPilA is located at the cell surface. To visualize CbPilA, we labeled *C. bescii* cells grown on xylan with anti-CbPilA polyclonal serum, and a fluorescently-tagged secondary antibody. We expect that the signal is specific for CbPilA and not the other predicted minor pilins, as the anti-CbPilA antibody minimally interacts with other predicted pilins at the dilution used for immunofluorescence microscopy (**Fig. S3**), and that previous proteomics studies did not detect any of the other minor pilins during growth on xylan (4, 21). A fluorescent signal from anti-CbPilA antibodies were observed surrounding *C. bescii* cells during growth on xylan (**Fig. 5B** **and C**), but not during growth on xylose (**Fig. E and F**). Based on the direct interaction of rCbPilA with xylan at 75°C (**Fig. 4**), we suspected that the larger fluorescing green clusters in **Fig. 5B**, were residual xylan particles coated with CbPilA. To verify this, we incubated xylan and rCbPilA at 75°C, and processed those samples for immunofluorescent microscopy to detect any CbPilA bound to xylan. As expected, we could detect a fluorescent signal from rCbPilA-treated xylan (**Fig. 5H**) which resembles the fluorescing green clusters in **Fig. 5B**. To ensure that the fluorescent signal was not the result of a non-specific interaction between anti-CbPilA and xylan, we included a biomass-only xylan control where uninoculated LOD medium was processed with both primary and secondary antibody (**Fig. S5H**). We were unable to detect a fluorescent signal, supporting that anti-CbPilA is not adsorbing to xylan, and the fluorescent signal detected with rCbPilA treated xylan (**Fig. 5H**) is a specific CbPilA-xylan interaction. Furthermore, *C. bescii* cells grown on xylan or xylose and processed with only the secondary antibody (**Fig. S5A-F**) lack any fluorescent signal, establishing that non-specific adsorption of the secondary antibody is not contributing to the signal.

**Figure 5:**
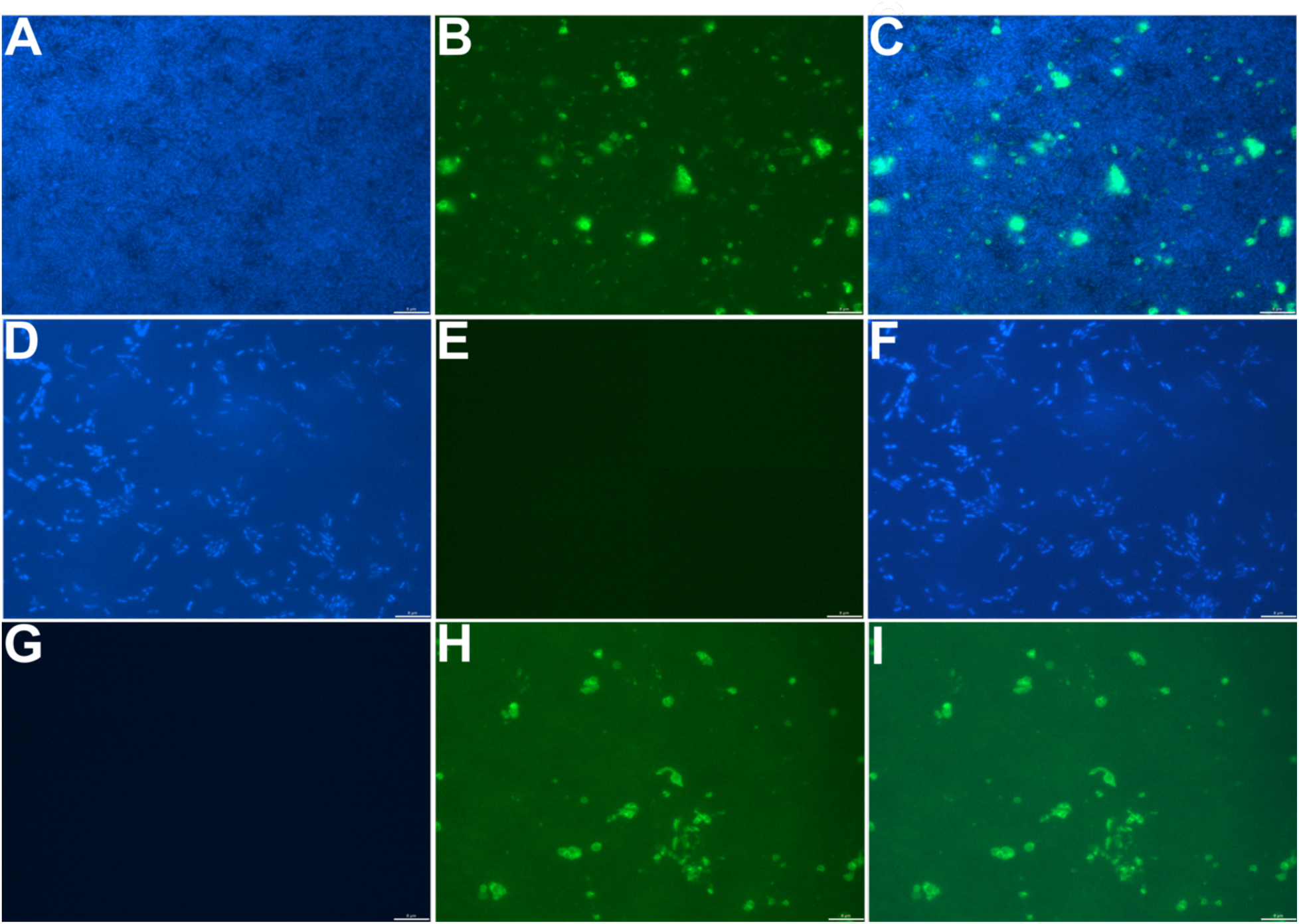
Immunofluorescence detection of extracellular CbPilA from *C. bescii* cultures. *C. bescii* cells (blue), or CbPilA (green) were visualized with epifluorescence microscopy. *C. bescii* cells grown on xylan (A, B, and C) or xylose (D, E, and F) were processed (A, B) DAPI, GFP and (C) color composite image of immunofluorescence microscopy of *C. bescii* cells grown on xylan. (D, E) DAPI, GFP and (F) color composite image of immunofluorescence microscopy of *C. bescii* cells grown on xylose. (G, H) DAPI, GFP and (I) color composite immunofluorescence microscopy of image of immunofluorescence microscopy of xylan treated with rCbPilA. Brightness and offset of DAPI (blue) and GFP (green) channels were balanced in the color composite images using Image-Pro Insight 9.1.

### <u>C. bescii</u> cell adhesion to polysaccharides is CbPilA dependent

Given the proximity of the T4P locus to tāpirin proteins and major cellulases used by *C. bescii* (10), we originally hypothesized that the major pilin was functioning as an adhesin, by binding to plant polysaccharides. Based on the affinity of rCbPilA for xylan (**Fig. 4, and** **Fig. 5**), we further examined its role in adherence by assessing if it would interfere with *C. bescii* attachment to insoluble polysaccharides, using a planktonic cell attachment assay (**Fig. S6**). Our experimental design allowed us to test whether the presence of rCbPilA and or substrate influenced the attachment of planktonic cells to insoluble substrates. In these assays, a reduction in planktonic cell densities (PCD) after treatment is indicative of cell attachment. Based on the increased production of CbPilA protein in response to xylan, we tested *C. bescii* cells grown on xylan for their ability to attach to insoluble xylan or cellulose (**Fig. 6A, C**). As a comparison, we also tested if cells grown on cellulose behaved similarly (**Fig. 6B**).

**Figure 6.**
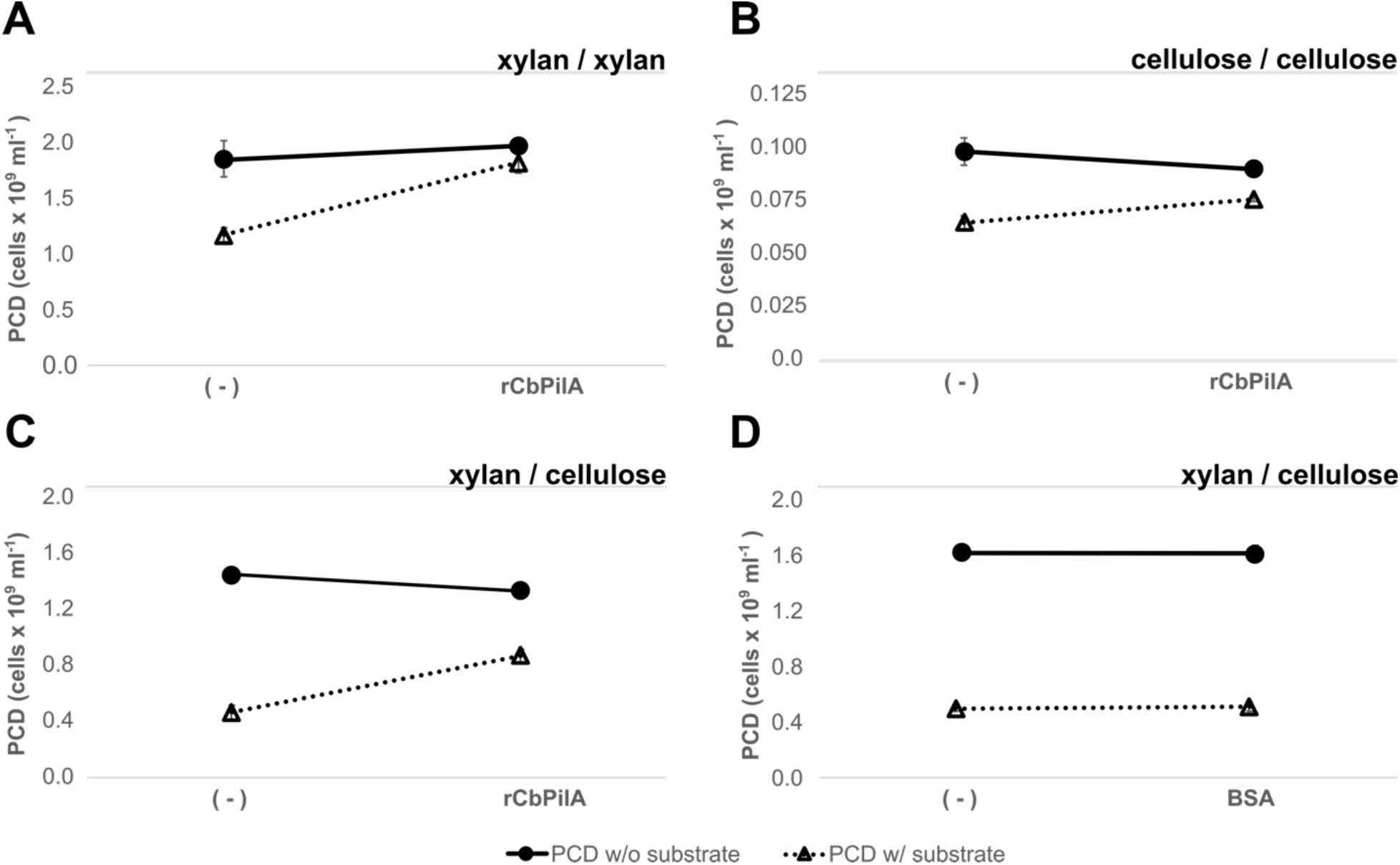
Cell binding assays showing inhibition effects of rCbPilA (or BSA) on binding of *C. bescii* cells to a xylan or cellulose substrate, in four separate experiments. Each panel displays planktonic cell densities as a function of two predictor variables: binding-substrate presence/absence and protein presence/absence (e.g. “(-)” versus “rCbPilA”). (A) *C. bescii* cells from cultures grown on xylan and incubated with xylan as a binding substrate. (B) *C. bescii* cells from cultures grown on cellulose and incubated with cellulose as a binding substrate. (C, D) *C. bescii* cells from cultures grown on xylan and incubated with cellulose as a binding substrate. Solid lines indicate planktonic cell densities (PCDs) without substrate whereas dotted lines indicate PCDs with substrate. “(-)”, PCD values when no rCbPilA or BSA was added; “rCbPilA”/”BSA”, either rCbPilA or BSA was added to the binding assay. Higher PCD values correlate with fewer cells attaching to substrate versus lower values of PCDs which correlate with more cells attaching to the substrate. A Two-way ANOVA revealed significant main and interactive effects of binding substrate and rCbPilA on average PCDs (all p-values < 0.02; Table S3), indicating that rCbPilA significantly altered the binding-substrate effect (i.e., binding affinity) in all three experiments. In contrast, there was no significant main or interactive effect of BSA on average PCD. Each experiment had at least 3 biological replicates. Error bars indicate standard error.

In all cases, the PCD after exposure to insoluble substrate (**Fig. 6**, dotted lines) were lower than the PCD of cells exposed to buffer alone (**Fig. 6**, solid line), indicating that *C. bescii* cells were attaching to xylan and cellulose, as expected. Surprisingly, while the attachment of *C. bescii* cells to xylan after growth on xylan (26% attachment, **Fig. 6A**) is expected, a higher proportion of cells were capable of attachment to cellulose after growth on xylan (69% attachment, **Fig. 6C**). Moreover, when cells are exposed to both rCbPilA and substrate (**Fig. 6A-C**) PCDs increase, indicating that the presence of rCbPilA during the attachment process is inhibiting the ability of cells to interact with the substrate. We observed a 3.3-fold decrease in cell attachment to xylan (**Fig. 6A**) compared to 2-fold decrease in cell attachment to cellulose (**Fig. 6C**). As expected, when *C. bescii* was incubated with substrates at elevated temperatures (75°C), we observed an increase in cells attached to xylan and cellulose compared to data collected at 25°C and with the addition of rCbPilA, cell attachment was decreased 6 and 24-fold for xylan and cellulose respectively (**Table S3**).

Two-way ANOVA was then used to test for interaction effects between our independent variables, substrate and protein (**Table S4**). The interaction between treatments (substrate and CbPilA) were statistically significant with p-values below 0.05 (**Table S4**). This interaction is also specific, as a control protein, BSA, did not interfere with attachment based on a lack of statistical interaction in this experiment (**Fig. 6D**, **Table S4**). As an additional test of specificity between *C. bescii* cells and polysaccharides, we tested the ability of two different strains of *E. coli* cultured in defined medium to adhere to xylan or cellulose (**Table S5**). In both cases, no significant decrease in *E. coli* PCD was observed, supporting the specificity of the cell attachment assay. By extension, our data support that *C. bescii* T4P play an integral role in cellular attachment to xylan, but additional mechanisms, such as the tāpirins, are involved in cellular attachment to crystalline cellulose.

To further confirm the role of *CbPilA* in attachment to plant polysaccharides like xylan and cellulose, we also tested if anti-CbPilA antibodies could neutralize CbPilA binding sites and interfere with the ability of planktonic cells to adhere to xylan or cellulose. Cells cultured on xylan were tested for their ability to adhere to xylan or cellulose when incubated with increasing concentrations of anti-CbPilA antibody serum. Pre-incubation of cells with anti-CbPilA antibodies was sufficient to interrupt cell adhesion, and resulted in up to 1.75-fold and 2.2-fold increases in PCD in the presence of xylan and cellulose, respectively (**Fig. 7**). These data are in line with data from cell binding assays (**Fig. 6**) and further support the role of *C. bescii* T4P in attachment to plant polysaccharides like xylan and cellulose.

**Figure 7.**
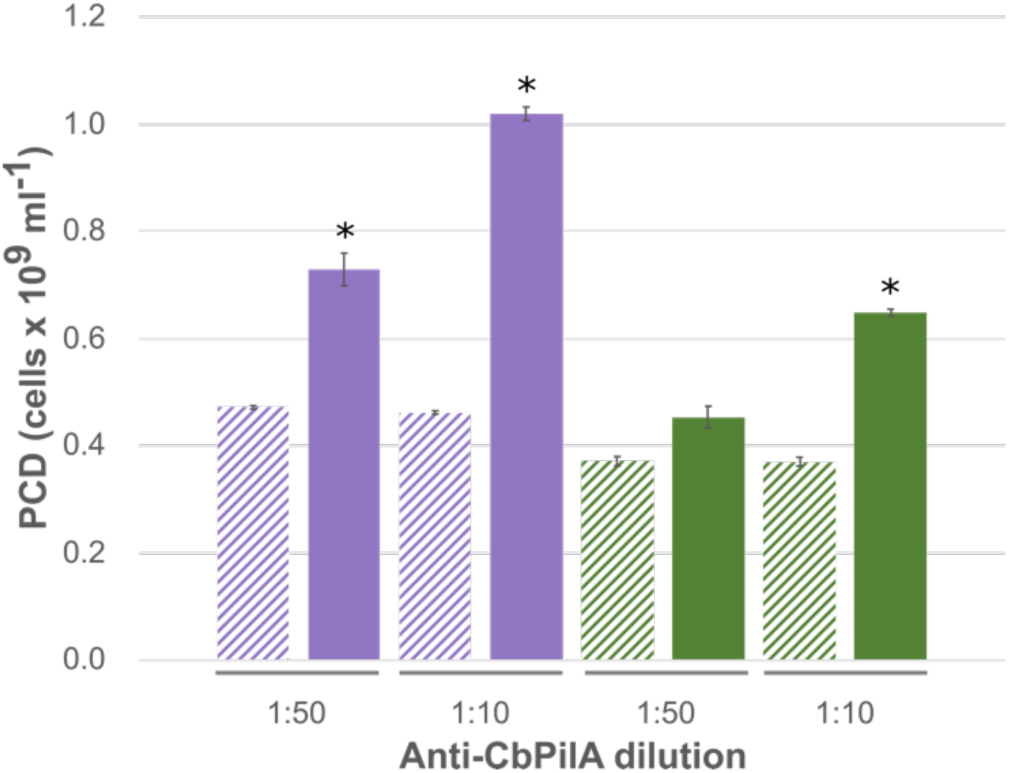
Pre-treatment with anti-CbPilA antibody increases planktonic cell density in the presence of insoluble polysaccharides. Planktonic cell densities (PCD) of *C. bescii* cells from grown on xylan were incubated with cellulose (purple bars) or xylan (green bars) as a binding substrate. Bars with diagonal lines indicate *C. bescii* cells pre-treated with pre-immune serum, solid bars are *C. bescii* cells pre-treated with anti-CbPilA antibodies. One way ANOVA was used to compare the average planktonic cell densities across all the treatment groups in the assay followed by a post hoc Tukey HSD test to check which groups had a statistically significant difference (*, p-value < 0.05). Error bars indicate standard error.

## DISCUSSION

In this study, we present an initial functional characterization of the Gram-negative like major pilin from *C. bescii*. Similar to other members of the class Clostridia (24, 29), the *C. bescii* genome encodes for the genes required to produce a Gram negative-like T4P in a single locus (**Fig. 1A**), that is likely arranged as an operon (**Table S1**). Initial bioinformatics analyses identified five putative pilin genes located in the *C. bescii* T4P locus (**Table 1**). Pathogenic Firmicutes, such as *Streptococcus sanguinis* (47, 48) and *C. perfringens* (28), also encode for multiple pilins in their respective T4P loci, and a hypervirulent *C. difficile* strain encodes for as many as nine pilins distributed across five genomic loci (30). Additionally, based on amino acid sequence homology, concatenated *C. bescii* pilin sequences clusters with other T4P loci from the majority of strongly cellulolytic *Caldicellulosiruptor* species (**Fig. 1B**).

Based on our phylogenetic analysis, and analysis of publicly available proteomics data (10), we hypothesized that the *C. bescii* T4P is likely used to facilitate cell attachment to polysaccharides found in plant biomass. To test this hypothesis, we first used transcriptomics and proteomics data (**Table 2**) to identify the *C. bescii* major pilin, (CbPilA) for molecular cloning and analysis. We sought to determine if members of the non-cellulosomal genus *Caldicellulosiruptor* used T4P to facilitate attachment to plant polysaccharides similar to observations in cellulosomal members of the genus *Ruminococcus* (26, 49). In comparison to *R*. *flavefaciens* or *F. succinogenes*, that produced pilin during growth on cellulose (23, 24) and/or glucose (23), we observed that the highest CbPilA production was observed during growth on xylan in comparison to glucomannan, xylose (**Fig. 3**), cellulose, or pectin (data not shown), implying that xylan is the main inducer of T4P locus expression in *C. bescii*. This is not completely unsurprising, considering that after cellulose, xylan is the second most common polysaccharide present in secondary plant cell walls (50). Response to soluble xylooligosaccharides in carbon-limited biotopes would be a likely adaptation to sense plant biomass in terrestrial hot springs. For example, *Hungateiclostridium thermocellum* (formerly ‘*Clostridium thermocellum’*) produces extracytoplasmic factor anti-sigma factors that upregulate certain cellulosome-related genes encoding for enzymes in response to soluble xylooligosaccharides in its environment (51, 52). Unlike in previous proteomics studies, we could not detect any extracellular CbPilA from cells grown on cellulose using immuno-dot blots, but this discrepancy with prior data may be explained by the use of complex media in the prior studies (4, 10, 35). In this study, we cultured *C. bescii* on a chemically defined medium, ensuring that the tested polysaccharide was the only available carbon source. The level of amino acid sequence diversity observed among pilins from the genus *Caldicellulosiruptor* (**Fig. 1B**) may also translate to differential genetic regulation of T4P loci. For example, highly cellulolytic species, *C. changbaiensis,* failed to adhere to xylan when used in cell binding assays (42), and likely deploys a T4P in response to other signals, or alternately, its pilin proteins are not involved in attachment to polysaccharides. *C. changbaiensis* was one of the highly divergent cellulolytic species (**Fig. 1B**) on the basis of pilin amino acid sequence, so it is not surprising that it would display different attachment properties in comparison to *C. bescii*.

In addition to identifying xylan as the inducing polysaccharide for CbPilA production, we demonstrated a direct interaction between rCbPilA and xylan *in vitro* with polysaccharide pulldown assays (**Fig. 4**) and immunofluorescence micrographs (**Fig. 5H**). As expected from a thermostable protein, we detected an interaction between rCbPilA and xylan at 75°C but not 25°C (**Fig. 4**), suggesting that physiological temperatures favor adsorption of CbPilA to carbohydrates, possibly through the exposure of hidden polysaccharide binding sites. However, planktonic *C. bescii* cells were capable of adhering to xylan and cellulose at 25°C and 75°C (**Table S3**) and anti-CbPilA antibody neutralization at 25°C was sufficient to interrupt *C. bescii* cell adhesion to xylan and cellulose (**Fig. 7**), indicating that elevated temperatures may only be required for recombinant forms of CbPilA to adhere to polysaccharides *in vitro*. Additionally, rCbPilA also interacts with cellulose (**Fig. 4**), similar to observations made using a pilin protein from *R. albus* 8 in a polysaccharide pull-down assay (25). By extension, these data support the role of the *C. bescii* T4P, via the major pilin CbPilA in cell attachment to plant polysaccharides.

Although we demonstrated a direct interaction between rCbPilA with xylan (**Figs. 4, and 5H, I**) and cellulose (**Fig. 4**), *in vitro*, our goal was to determine whether CbPilA played a direct role in *C. bescii* cell attachment to these polysaccharides. The ability of *C. bescii* cells to attach to xylan or cellulose were monitored using planktonic cell attachment assays to insoluble polysaccharides (**Figs. 6**, **7** **and Table S3**). This attachment was, in part, mediated by T4P in a specific manner, as we observed a decrease in adhered cells to xylan or cellulose after supplementation with rCbPilA (**Fig. 6**) or anti-CbPilA antibody neutralization (**Fig. 7**). It is likely that the observed inhibition of cell attachment to xylan (**Fig. 6A**) is a result of rCbPilA binding to xylan, as observed in **Fig. 6I**, and blocking available binding sites for native CbPilA. Overall, our observations from these planktonic cell attachment assays support the role of CbPilA and by extension it’s T4P in attachment to plant polysaccharides. T4P mediated attachment to the plant polysaccharide, cellulose was previously demonstrated by members of the genus *Ruminococcus* (26, 49).

Taken together, our *in vitro* and *in vivo* data provides evidence that CbPilA plays a direct role in *C. bescii* adherence to xylan. While other members of the genus *Caldicellulosiruptor* produce an S-layer located xylanase that facilitates cell attachment to xylan (9), *C. bescii* lacks this gene, and instead deploys type IV pili to adhere to xylan (**Fig. 8**). In addition, CbPilA may function along with the tāpirins (19, 22) to promote cell adherence to cellulose and we propose a model where cell adhesion to cellulose involves both proteins (**Fig. 8**). A similar mechanism has been recently described for a T4b pilus from enterotoxicogenic *Escherichia coli*, where a secreted adhesin (CofJ) protein functions as a type of “scout” that first associates with receptors on the host cell, and subsequently interacts with minor pilins present at the tip of the T4b pilus (53). Future studies to confirm the interaction of tāpirin proteins with T4 pilins from the genus *Caldicellulosiruptor* are underway. Overall, our data support the idea that non-cellulosomal species, like those from the genus *Caldicellulosiruptor*, have evolved alternate mechanisms to adhere to insoluble substrates.

**Figure 8.**
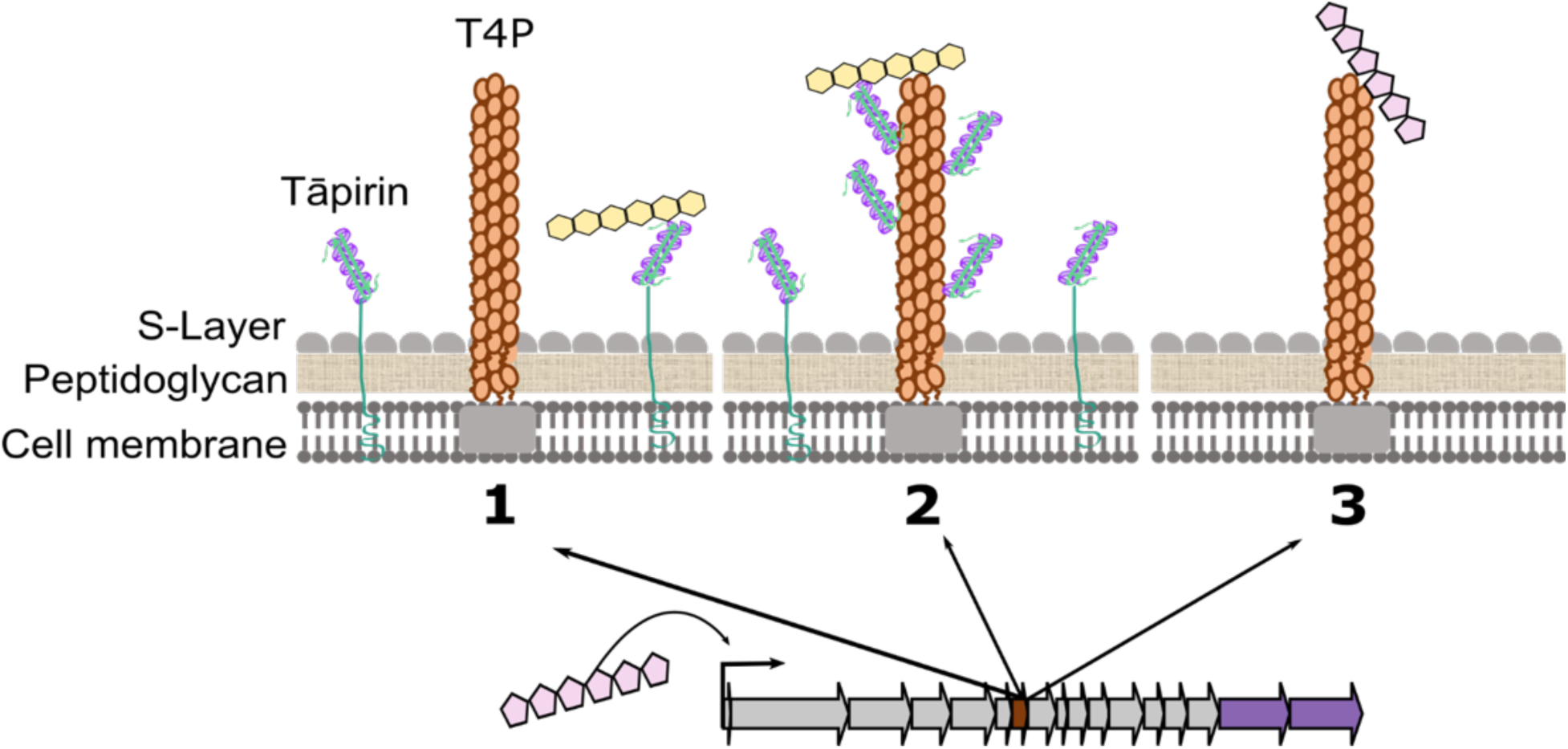
Model of the potential role(s) of type IV pili during deconstruction of plant biomass by C. bescii. Imported xylooligosaccharides induce expression of the type IV pilus operon (brown arrow, CbPilA; purple arrows, tāpirins). 1, Tāpirins interact directly with cellulose. 2, Type IV pili interact with cellulose in association with tāpirins. 3, Type IV pili interact directly with xylan. Cream colored hexagons, cellooligosaccharides; pink pentagons, xylooligosaccharides.

## MATERIALS AND METHODS

### Identification of Pilin Genes from *C. bescii* and bioinformatics analysis

Putative pilin genes in *C. bescii* were identified using the Joint Genome Institute Integrated Microbial Genomes (IMG) database (54). Functional annotation of the predicted amino acid sequences for each pilin used the Pfam database (55) available within IMG. Signal peptides were predicted by the SignalP 4.1 Server (56) and transmembrane domains were predicted using TMHMM, version 2.0 (57). Hairpin sequences that may serve as transcriptional terminators were identified using TransTermHP (58). BLASTp, hosted within the IMG database, was used to identify *Caldicellulosiruptor* orthologs to the five pilins encoded in the *C. bescii* T4P operon (54, 59). Amino acid sequences of orthologs were aligned using MUSCLE (60) hosted on the EMBL-EBI server (61). Sequence Matrix (https://github.com/gaurav/taxondna) was used to concatenate the amino acid alignments prior to the generation of a phylogenetic tree. IQ-TREE (62) along with substitution model selection by ModelFinder (63) was used to generate a maximum-likelihood phylogenetic tree, which was tested by 1000 bootstraps using UFBoot (64), all of which were hosted by the Los Alamos National Laboratory (https://www.hiv.lanl.gov/content/sequence/IQTREE/iqtree.html). Secondary structures of putative *C. bescii* pilins were predicted using the PSIPRED program, version 3.3 (65).

### Media, growth conditions and estimation of cell densities

Minimal, low osmolality defined (LOD) medium (66) was used for culturing *C. bescii* DSM 6725 strains on various carbon sources. *C. bescii* DSM 6725 was provided by Robert M. Kelly (North Carolina State University, Raleigh, NC). *C. bescii* cultures were grown at 75°C under anaerobic conditions with one of the following substrates, all at 1 g l^-1^: xylose, Sigmacell (20 µm, Sigma), xylan (beechwood, Sigma), glucomannan (konjac root, NOW Foods). Epifluorescence microscopy (Nikon Eclipse E400) was used to enumerate glutaraldehyde fixed cells stained with acridine orange as previously described (67).

*Escherichia coli* was maintained on LB medium supplemented with antibiotics (25 mg l^-1^ kanamycin, 34 mg l^-1^ chloramphenicol) for plasmid maintenance or ZYM-5052 autoinduction medium supplemented with 50 mg l^-1^ and 34 mg l^-1^ (68) for protein production. For cell binding assays, *E. coli* was cultured on glucose M9 minimal medium (6 g l^-1^ Na_2_HPO_4_, 3 g l^-1^ KH_2_PO_4_, 0.5 g l^-1^ NaCl, 1 g l^-1^ NH_4_Cl, 1mM MgSO_4_•7H_2_O, 0.1mM CaCl_2_, and 2 g l^-1^ glucose). Cell densities of *E. coli* cultures were measured as optical density at 600 nm (OD_600_) by a spectrophotometer.

### Production and purification of recombinant proteins

Soluble protein was produced from truncated *cbpilA*, (nucleotides 112 – 438, GenBank accession number WP_015908263.1). Molecular cloning into pET-28b used restriction enzyme sites NcoI and XhoI incorporated into oligonucleotide primer sequences, respectively (Forward primer, CGC<u>CATATG</u>CAAGTATTAAAACAGATAAAC; Reverse primer, GCTA<u>CTCGAG</u>TTACTTGTAGTTTGGATCTGG). Soluble rCbPilA was produced from a truncation mutant that eliminated the pre pilin-type N-terminal cleavage sequence and a region encoding for the α-1N helix (see **Fig. S2**). All minor pilins were cloned into pET-28b using Gibson Assembly, per manufacturer’s protocols (NEB). Soluble recombinant *C. bescii* minor pilins, Athe_1881 (GenBank accession *WP_015908264.1,* Forward primer: CTGGTGCCGCGCGGCAGCCAATTAAAGAATATAAATAAGGCAAGAAAG and Reverse primer: AGTGGTGGTGGTGGTGGTGCTTAGTAACCAGGATCAACATTTG); Athe_1877 (GenBank accession *WP_015908260.1*), Athe_1876 (GenBank accession *WP_015908259.1*) and recombinant *C. bescii* tāpirin (rCbtāpirin), Athe_1871 (GenBank accession YP_002573732 Forward primer: CTGGTGCCGCGCGGCAGCCATATGTTAGCATCGCTGAACCAG and reverse primer: AGTGGTGGTGGTGGTGGTGCTCGAGCTACCTTGTAACCATGTTTC) were cloned, produced purified and dialyzed similar to rCbPilA. All recombinant pilins were produced with an *N*-terminal histidine tag for immobilized nickel affinity purification using 1ml HisTrap columns (GE Healthcare) per the manufacturer’s protocol. Auto-induction medium (68) supplemented with 50 mg l^-1^ kanamycin and 34 mg l^-1^ chloramphenicol was used for growth of *Escherichia coli* Rosetta (Novagen) to induce protein production. Purity of recombinant pilin proteins were confirmed by SDS-PAGE after dialysis against 50mM sodium phosphate (pH 7.4) using SnakeSkin dialysis tubing (Thermo-Scientific) per the manufacturer’s protocol. The final concentration of recombinant *C. bescii* pilins was determined using the bicinchoninic acid assay (BCA, Thermo Scientific) per the manufacturer’s protocol.

### Western blots

rCbPilA and *C. bescii* minor pilins Athe_1881, Athe_1877, Athe_1876 were purified as described above and the histidine tags were cleaved off using thrombin. For thrombin cleavage, all proteins were dialyzed in thrombin cleavage buffer (tris-HCl, 50mM, NaCl 20mM, pH 8.0) and incubated with 15µl of the thrombin beads per mg of the respective protein for 7 hours at room temperature followed by overnight at 4°C with gentle shaking on an orbital shaker for the entire incubation period. Complete cleavage of the proteins was confirmed using SDS-PAGE and purified uncleaved recombinant pilins as well as thrombin cleaved recombinant proteins were further analyzed using western blotting and detection with an anti-CbPilA antibody. Heat-treated cell lysate from *E. coli* transformed with pET-28b(+) was used as a negative control. All protein fractions were equalized to a concentration of 30 µM, resolved on an SDS-PAGE gel as described previously. Proteins were transferred from the SDS-PAGE gel on to a PVDF membrane (5.5 cm x 8.5 cm) in a transfer cassette at 20V overnight at 4°C. Thereafter the membrane was probed and imaged as described for the immuno-dot-blots below. 30 µM of each thrombin cleaved pilin was also analyzed by on separate immuno-dot blots with antibody dilutions used for immuno-dot blots (for monitoring CbPilA production) described below as well as for immunofluorescence assays.

### Immuno-dot blots

Actively growing cultures of *C. bescii* on plant polysaccharides xylan, glucomannan, cellulose and the sugar xylose were sampled throughout growth until cultures reached stationary phase. Planktonic cells in growth medium were sampled at early, mid and late exponential growth diluted 1:1 with sterile glycerol prior to storage at -20° C until further use. These cell samples were used in the immuno-dot blots without any additional processing. Purified rCbPilA was applied on each membrane as a standard curve for quantitative immunoblot analysis. PVDF membranes used for protein immobilization (Amersham Hybond 0.2 PVDF, GE) were pre-wet with 100% methanol per the manufacturer’s instructions and then equilibrated in 1X PBS buffer. Samples were applied to the pre-wetted membrane blocked in PBS-T blocking buffer (PBS with 0.1% Tween-20 (v/v), 5% milk (w/v)) for one hour on an orbital shaker at room temperature. Afterwards, the membrane was rinsed with PBS-T washing buffer. Primary antibody (chicken anti-CbPilA, GeneTel) was diluted 1:1000 in PBST and incubated at room temperature. Afterwards, the membrane was washed in PBS-T. Secondary antibody (rabbit anti-chicken HPR conjugate, Immunoreagents) was diluted 1:1000 PBS-T and again incubated at room temperature. The membrane was then washed in PBS-T as previously described. Chemiluminescent detection of secondary antibody used ECL western blotting detection reagents (GE), per the manufacturer’s protocol. Membranes were imaged (ChemiDoc Touch Imaging System, Bio-Rad) using standard chemiluminescent settings and analyzed using Image Lab 5.2.1 (Bio-Rad). Three independent replicates were blotted for each time point. Cell densities determined while plotting growth curves were used to calculate the number of cells applied to the membrane at each time point. A linear relation between the signal intensity and concentration of CbPilA in the sample loaded was verified by a standard curve (R-squared = 0.98). Standards included purified rCbPilA (1 ng µl^-1^, 10 ng µl^-1^, 50 ng µl^-1^, 100 ng µl^-1^ and 200 ng µl^-1^). Native CbPilA concentrations in the samples were determined using rCbPilA standards and were then normalized to the number of cells, and compared using a two-sample t-test in R studio statistical software (69) (v.3.3.3).

### Immunofluorescence microscopy

Immunolabeling of *C. bescii* cells followed methods from Conway et al. (9) with modifications. *C. bescii* cells were grown on 50 mL LOD medium in 125 ml serum bottles with either xylan or xylose as the carbon source and harvested at late exponential phase. Residual xylan was separated from the cells and cells were subsequently pelleted using centrifugation 5000 x g for 10 minutes at room temperature (same conditions for all of the centrifugation steps). Anti-CbPilA polyclonal antibodies were raised in chicken (chicken anti-CbPilA, GeneTel), and used as the primary antibody at a 1:120 dilution. The secondary antibody was a goat, anti-chicken IgY polyclonal antibody conjugated to DyLight 488 (Novus Biologicals) used at a 1:400 dilution. After washing the primary antibody, CbPilA was labeled with the secondary antibody at room temperature for an hour followed by staining of cells were with 1µg/ ml DAPI in 1X PBS for 5 minutes at 4°C. After the final washing step, cells were then vacuum-filtered onto 0.2µm polycarbonate track etched membrane filters (GVS Life Sciences) and mounted in Vectashield mounting medium for imaging (Vector laboratories). Epifluorescence imaging used a Nikon eclipse E400 microscope and images were captured using an Infinity 3 Lumenera camera. DAPI and GFP images were merged into a color composite image using Image-Pro Insight software 9.1 (Media Cybernetics). Secondary control images were captured from secondary antibody labeled samples which were not incubated with the primary antibody. Biomass controls (uninoculated xylan in LOD) were processed with both primary and secondary antibodies. Xylan was incubated with 30 µM rCbPilA at 75°C for an hour similar to the polysaccharide pull down assay and then probed with primary and secondary antibodies and imaged as the other samples. Brightness and offset of DAPI (blue) GFP (green) channels were balanced in the color composite images using Image-Pro Insight 9.1. All assays had biological triplicates. Scale bars at the bottom of each image indicate 8µm.

### Polysaccharide pull down assays

Substrates (cellulose or xylan) used in all polysaccharide pull down assays were washed with distilled water five times followed by 16h air drying and then at 70°C for two hours. Substrate (100 mg ml^-1^) was incubated with rCbPilA (30 µM) in the binding buffer, 50 mM sodium phosphate, pH 7.2 and incubated in a thermomixer (Eppendorf) at 700 rpm for one hour at 25°C or 75°C. Insoluble substrates were pelleted by centrifugation for one minute (15000 x g). The supernatant (∼ 70µl) represented unbound protein. Pelleted substrate was washed with 1 ml of the binding buffer five times. After the final wash, the pellet was resuspended in 70µl of binding buffer representing the bound protein. Both the bound and unbound fractions were analyzed using SDS-PAGE.

### Cell binding assays

*C. bescii* cell cultures were grown to early stationary phase supplemented with either xylan or cellulose (1 g l^-1^) and harvested at 5000 x g for 10 minutes at room temperature. Cells were resuspended in 50 mM sodium phosphate, pH 7.2 to a density of 109 cells ml^-1^. Each experimental condition consisted of a total volume of 1.2 ml comprised of 1 ml *C. bescii* cells, and 0.2 ml of either rCbPilA (30 µM), BSA (30 µM) or buffer. Washed xylan or cellulose (100 mg ml^-1^, as described above) was added to the experimental conditions, and no binding substrate was added to the negative controls. All binding assays were incubated at room temperature or 75°C with gentle rotary shaking at 100 rpm for one hour. After incubation, planktonic cells were enumerated using epifluorescence microscopy as described above. Each of the four treatments in all cell binding assays were replicated a minimum of three times. Results from each experiment were analyzed with a two-way ANOVA, using the functions “lm” and “Anova” in Program R (70). Each model tested for main and interactive effects of binding substrate (presence/absence of xylan or cellulose) and protein (presence/absence of rCbPilA or BSA) on the planktonic cell density. In this modeling framework, the significance of the interaction term (Substrate×Protein) indicates whether the presence of the protein (rCbPilA or BSA) influenced the binding affinity of cells to the substrate (xylan or cellulose).

As an additional negative control, *E. coli* strains NEB10β or BL21 were cultured overnight in glucose M9 minimal medium (6 g l^-1^ Na^2^HPO_4_, 3 g l^-1^ KH_2_PO_4_, 0.5 g l^-1^ NaCl, 1 g l^-1^ NH_4_Cl, 1mM MgSO_4_•7H_2_O, 0.1mM CaCl_2_, and 2 g l^-1^ glucose). Cells were harvested as described above for *C. bescii*, washed once in binding buffer, 50 mM sodium phosphate, pH 7.2, and resuspended to an approximate OD_600_ of 1.0. Cell binding to washed xylan or cellulose followed the same protocol as for *C. bescii*, with two exceptions, no rCbPilA was used as an experimental condition, and planktonic cell densities were measured in 96-well polystyrene plates using a microplate reader (Biotek Epoch). Each experimental condition was replicated a minimum of three times.

### Cell binding assays with antibody neutralization

Antibody neutralization assays followed Mahmoud *et a*l (71) with modifications. rCbPilA was replaced with dilutions of polyclonal anti-CbPilA. Cells were grown and harvested as described above, and incubated with a 1:50 or 1:10 dilution of anti-CbPilA fpor experimental conditions, and similar dilutions of pre-immune chicken serum served as the negative control. Both control and treated cells were incubated with and without insoluble xylan or cellulose and enumerated using epifluorescence microscopy as described above. Each treatment in all cell binding assays had three biological replicates. Average planktonic cell densities for each treatment were compared using a one-way ANOVA, using the functions “lm” and “Anova” in Program R (70) followed by post hoc Tukey HSD tests using R software (70).

## ACKNOWLEDGEMENTS

The authors are grateful for the technical assistance by Monique Clemmon and Leandra Egner in molecular cloning of the minor pilins. This study was supported by start-up funds to S.E.B.-S. (Oakland University). A.K. was supported in part through funds from the Center for Biomedical Research (OU).

## SUPPLEMENTARY DATA

Table S1. Hairpin sequences present in the type IV pilus locus

Table S2. **Percent identity (amino acid) of each CbPilA homolog across 14 sequenced *Caldicellulosiruptor* genomes**

**Table S3.** Effect of rCbPilA on planktonic *C. bescii* cell densities after incubation with xylan or cellulose at 75°C

**Table S4. Cell attachment analysis of variance results**

**Table S5.** Planktonic *E. coli* cell densities after incubation with xylan or cellulose

**Figure S1. Alignment of *Caldicellulosiruptor bescii* hypothetical proteins with putative pilin leader sequences.** The conserved glycine (G, purple) and glutamic acid (E, green) from the pre-pilin N-terminus cleavage domain (GFxxxE) are highlighted.

**Figure S2. Cloning strategy to produce soluble rCbPilA. A** truncation mutant of Athe_1880 was designed to remove the prepilin-type N-terminal cleavage/methylation domain-GFxxxE (cyan hexagon), as well as the highly hydrophobic region (magenta) and the predicted hydrophobic transmembrane domain (green). Ruler units are in base pairs.

**Figure S3. Immuno-analysis of *C. bescii* pilins with anti-CbPilA.** (A) Immunoblot analysis of *C. bescii* pilins with anti-CbPilA. rCbPilA, Athe_1881, Athe_1877 and Athe_1876 were purified as described in the methods and the 6xHistags were cleaved using thrombin. Uncleaved as well as cleaved proteins were quantified, equalized to the same molar concentration (30 µM) and an equal volume of these pilins was loaded in each well. Heat treated *E. coli* cell lysate carrying an empty pET-28b expression vector was used as a negative control. (B) Immuno-dot blots probed using a 1:120 primary antibody dilution used for immunofluorescent microscopy. (C) Immuno-dot blots probed using an 1:1000 primary antibody dilution used for immuno-dot blots to quantify CbPilA expression. (D) Densitometry analysis from the immuno-dot blots in panel *B*. (E) Densitometry analysis for immuno-dot blots in panel *C*. For B and C, rCbPilA, Athe_1881, Athe_1877 and Athe_1876 were purified as described in the methods and 6xHis tags were cleaved using thrombin. Uncleaved as well as cleaved *C. bescii* pilins were quantified, equalized to the same molar concentration (30 µM) and an equal volume of these pilins was spotted on a PVDF membrane. For D and E, each column represents average intensity of the 5 replicates of each sample. Error bars represent standard error.

**Figure S4. Validation of the polysaccharide pulldown assay with recombinant *C. bescii* tāpirin.** (A) SDS-PAGE analysis of bound and unbound fractions of rCbtāpirin. 1, bound fraction of rCbtāpirin; 2, unbound fraction of rCbtāpirin. Polysaccharides tested were xylan and microcrystalline cellulose (Sigmacell). Negative control includes only rCbtāpirin without the polysaccharide. SDS-PAGE images are representatives of the results from assays run in triplicate. (B) Densitometry analysis for rCbtāpirin bound to cellulose at 75°C (C) Densitometry analysis for rCbtāpirin bound to xylan at 75°C. For B and C, each column represents average normalized intensity of the 3 replicates of each respective assay. Error bars represent standard error.

**Figure S5. Secondary antibody controls and biomass control for immunofluorescence detection of extracellular CbPilA from *C. bescii* cultures.** (A, B) DAPI, GFP and (C) color composite image of immunofluorescence microscopy of *C. bescii* cells grown on xylan and processed only with the secondary antibody. (D, E) DAPI, GFP and (F) color composite image of immunofluorescence microscopy of *C. bescii* cells grown on xylose and processed only with the secondary antibody. (G, H) DAPI, GFP and (I) color composite immunofluorescence microscopy of image of immunofluorescence microscopy of uninnoculated LOD xylan medium processed with both primary and secondary antibodies. Brightness and offset of DAPI (blue) GFP (green) channels were balanced in the color composite images using Image-Pro Insight 9.1. Scale bars at the bottom of each image indicate 8µm.

**Figure S6. Experimental design for the *C. bescii* cell binding assay.** Blue rod: *C. bescii* cell, green circle: substrate. Four different treatment groups were set up to test the effect of two independent variables-substrate and protein. For the substrate variable the two treatments levels were: cells with binding buffer (50 mM sodium phosphate pH 7.2) alone and cells with binding buffer and substrate. For the protein variable the two treatment levels were, cells with rCbPilA alone and cells with rCbPilA and substrate.

