## Supplementary Data for "*Caldicellulosiruptor bescii* regulates pilus expression in response to the polysaccharide, xylan"


---

### Supplementary Tables and Figures


**Table S1.** Hairpin sequences present in the type IV pilus locus

**Table S2.** Percent identity (amino acid) of each CbPilA homolog across 14 sequenced *Caldicellulosiruptor* genomes

**Table S3.** Cell attachment analysis of variance results

**Table S4.** Planktonic *E. coli* cell densities after incubation with xylan or cellulose

**Figure S1.** Alignment of *Caldicellulosiruptor bescii* hypothetical proteins with putative pilin leader sequences

**Figure S2.** Cloning strategy to produce soluble rCbPilA

**Figure S3.** Immuno-analysis of *C. bescii* pilins with anti-CbPilA

**Figure S4:** Validation of the polysaccharide pulldown assay with recombinant *C. bescii* tāpirin

**Figure S5.** Secondary antibody controls and biomass control for immunofluorescence detection of extracellular CbPilA from *C. bescii* cultures

**Figure S6.** Experimental design for the *C. bescii* cell binding assay

**Table S1. Hairpin sequences present in the type IV pilus locus**

| Upstream Gene Locus | Terminator Sequence | TTHP[a] Score | Predicted/ Confirmed | Note | Reference |
|---|---|---|---|---|---|
| Athe_1880 | CAATTTGGACAATGA **CTCGAGGGGAAT** ATAATTG **ATTCCCCTCGAG** GTATTCTGTATATGC | 100 | Predicted | Internal to Athe_1879 | This study |
| Athe_1871 | AATGGATATAAAAAA **GGCATCTGAC** GGTT **GCCAGATGCC** ATTAGCCAAGGATTA | 100 | Confirmed | No termination | (1) |
| Athe_1870 | AGCCACTTGCAAAAA **CACTCA** AAA **TGAGCG** TTTCCACACTCTTTA | 76 | Confirmed | Termination | (1) |

[a] TTHP, TransTermHP (http://transterm.ccb.jhu.edu/)

**Table S2. Percent identity (amino acid) of each CbPilA homolog across 14 sequenced *Caldicellulosiruptor* genomes.**

| | Calac[a] | Cbes | Calcha | Calda | Calhy | Calkr | Calkro | Calla | Calmo | Calna | COB47 | Calow | Csac | F32 | MAG |
|---|---|---|---|---|---|---|---|---|---|---|---|---|---|---|---|
| **Calac** | 100 | 54.23 | 72.79 | 54.23 | 50 | 53.64 | 54.23 | 56.74 | 41.79 | 53.9 | 52.11 | 53.24 | 52.03 | 54.23 | 53.19 |
| **Cbes** | 54.23 | 100 | 55.56 | 92.47 | 45.4 | 71.9 | 92.41 | 72.6 | 42.5 | 93.1 | 91.72 | 42.95 | 87.59 | 88.28 | 91.72 |
| **Calcha** | 72.79 | 55.56 | 100 | 55.94 | 47.62 | 54.61 | 56.25 | 56.03 | 37.5 | 56.25 | 55.56 | 47.65 | 53.47 | 54.17 | 56.25 |
| **Calda** | 53.5 | 92.47 | 55.94 | 100 | 44.08 | 71.92 | 91.1 | 72.6 | 42.54 | 90.4 | 87.67 | 44.67 | 86.99 | 86.99 | 87.67 |
| **Calhy** | 50 | 45.39 | 48.25 | 44.08 | 100 | 49.3 | 44.74 | 47.97 | 31.94 | 44 | 42.67 | 42.54 | 44.74 | 44.08 | 43.33 |
| **Calkr** | 55.4 | 71.92 | 54.61 | 71.92 | 47.9 | 100 | 71.23 | 93.06 | 42.54 | 71.23 | 71.92 | 45.52 | 70.55 | 71.23 | 71.92 |
| **Calkro** | 54.23 | 92.41 | 56.25 | 91.1 | 44.74 | 71.23 | 100 | 71.92 | 41.05 | 90.34 | 89.66 | 42.28 | 91.72 | 93.1 | 89.66 |
| **Calla** | 56.74 | 72.6 | 56.03 | 72.6 | 47.9 | 93.06 | 71.92 | 100 | 41.05 | 73.3 | 71.92 | 45.14 | 70.55 | 71.23 | 71.92 |
| **Calmo** | 41.79 | 42.5 | 32.06 | 42.22 | 32.61 | 42.54 | 41.05 | 41.04 | 100 | 41.8 | 42.54 | 35.13 | 40.88 | 40.88 | 41.5 |
| **Calna** | 53.9 | 93.1 | 56.25 | 90.41 | 44 | 71.23 | 90.34 | 73.29 | 41.79 | 100 | 92.41 | 42.95 | 85.52 | 86.21 | 91.72 |
| **COB47** | 52.48 | 91.7 | 55.56 | 87.67 | 42.6 | 71.92 | 89.66 | 71.92 | 41.79 | 92.4 | 100 | 42.95 | 87.59 | 89.66 | 99.31 |
| **Calow** | 53.24 | 42.95 | 47.65 | 44.74 | 42.54 | 45.52 | 42.28 | 46.21 | 35.51 | 42.95 | 42.95 | 100 | 43.28 | 42.76 | 42.28 |
| **Csac** | 53.52 | 87.59 | 53.47 | 86.99 | 44.74 | 70.55 | 91.72 | 70.55 | 40.88 | 85.5 | 87.59 | 44.3 | 100 | 97.24 | 87.59 |
| **F32** | 54.23 | 88.28 | 54.17 | 86.99 | 44.08 | 71.23 | 93.1 | 71.23 | 40.88 | 86.21 | 89.66 | 42.76 | 97.24 | 100 | 89.66 |
| **MAG** | 52.82 | 91.72 | 56.25 | 87.67 | 43.33 | 71.92 | 89.66 | 71.92 | 41.5 | 91.72 | 99.31 | 42.28 | 87.59 | 89.66 | 100 |

[a] *Caldicellulosiruptor* species names abbreviations and GenBank accession numbers: Calac, *C. acetigenus* (WP_029227959.1); Cbes, *C. bescii* (WP_029227959.1); Calda, *C. danielli* (WP_029227959.1); Calhy, *C. hydrothermalis* (WP_029227959.1); Calkr, *C. kristjansssonii* (WP_029227959.1); Calkro, *C. kronotskyensis* (WP_029227959.1); Calla, *C. lactoaceticus* (WP_029227959.1); Calmo, *C. morganii* (WP_029227959.1); Calna, *C. naganoensis* (WP_029227959.1); COB47, *C. obsidiansis* (WP_029227959.1); Calow, *C. owensensis* (WP_083792173.1) ; Csac, *C.saccharolyticus* (WP_083792173.1); CF32, *Caldicellulosiruptor* str. F32 ( WP_039767876.1); Calcha, *C. changbaiensis* (WP_127352241.1) Column, query.

| **Table S3. Effect of rCbPilA on planktonic *C. bescii* cell densities after incubation with xylan or cellulose at 75°C** | | | |
|---|---|---|---|
| **Buffer** | | **rCbPilA** | |
| **Control**[a] | **Xylan**[b] | **Control**[c] | **Xylan**[d] |
| 9.50E+08 ± 5.25E+06 | 6.01E+08 ± 1.05E+07 | 8.75E+08 ± 1.97E+07 | 8.23E+08 ± 9.19E+06 |
| **Control**[a] | **Cellulose**[e] | **Control**[c] | **Cellulose**[f] |
| 1.35E+09 ± 3.92E+08 | 4.24E+08 ± 4.07E+07 | 8.65E+08 ± 3.28E+07 | 8.41E+08 ± 2.49E+07 |

[a] Average planktonic cell density calculated after incubation in buffer only
[b] Average planktonic cell density calculated after incubation with buffer and xylan
[c] Average planktonic cell density calculated after incubation in rCbPilA only
[d] Average planktonic cell density calculated after incubation with rCbPilA and xylan
[e] Average planktonic cell density calculated after incubation with buffer and microcrystalline cellulose
[f] Average planktonic cell density calculated after incubation with rCBPilA and microcrystalline cellulose

| Table S4. Cell attachment analysis of variance results | | | | | |
|---|---|---|---|---|---|
| Experimental Conditions [a] | | | | | |
| Carbon Source | Bound To | Predictors | d.f.[c] | F value | p-value |
| Xylan | Xylan | Xylan | 1, 20 | 16.42 | 0.0006 |
| | | PilA | 1, 20 | 14.57 | 0.001 |
| | | Xylan × PilA[b] | 1, 20 | 6.64 | 0.017 |
| Microcrystalline Cellulose | Microcrystalline Cellulose | Cellulose | 1, 8 | 165.2 | < 0.0001 |
| | | PilA | 1, 8 | 24.3 | 0.0011 |
| | | Cellulose × PilA[b] | 1, 8 | 20.8 | 0.0018 |
| Xylan | Microcrystalline Cellulose | Cellulose | 1, 32 | 287.9 | < 0.0001 |
| | | PilA | 1, 32 | 11.03 | 0.0022 |
| | | Cellulose × PilA[b] | 1, 32 | 37.62 | < 0.0001 |
| Xylan | Microcrystalline Cellulose | Cellulose | 1, 16 | 809.1 | < 0.0001 |
| | | BSA | 1, 16 | 0.026 | 0.87 |
| | | Cellulose × BSA[b] | 1, 16 | 0.046 | 0.83 |

[a] Response variable: Planktonic cell density; predictor variables: Binding substrate (xylan or cellulose) and Protein (PilA or BSA)

[b] Interaction between the two predictor variables in each model, indicating whether the binding-substrate effect (binding affinity) is influenced by the presence of the Protein (PilA or BSA)

[c] Degrees of freedom

| Table S5. Planktonic *E. coli* cell densities after incubation with xylan or cellulose | | | |
|---|---|---|---|
| | **Control**[a] | **Xylan**[b] | **Cellulose**[c] |
| *E. coli* BL21 | 0.777 ± 0.020 | 0.798 ± 0.010 | 0.779 ± 0.028 |
| *E. coli* NEB10β | 0.669 ± 0.013 | 0.693 ± 0.015 | 0.676 ± 0.008 |

[a] Average planktonic cell density measured as $OD_{600}$ after incubation in buffer only
[b] Average planktonic cell density measured as $OD_{600}$ after incubation with xylan
[c] Average planktonic cell density measured as $OD_{600}$ after incubation with microcrystalline cellulose

```
                 -20          -10         -1 1   5     10            20           30
                  |            |           | |   |     |             |            |  Leader Sequence
Athe_1872                          MKLKG FSLIEAIVAV AIFAIILVPI GIAFNQSIRV          5
Athe_1876               MRFNLEIKSITKLKG LTLVELIIAV VIVAIILGAI YSFFIQNFKV         15
Athe_1877                          MKCRG FSLSEMIIVI VIIAILIAIG IPAYLSSVRR          5
Athe_1880               MMAWLVKQVNSKHKG FTLIEMVIVV AIIAILIAIA VPQVLKQINK         15
Athe_1881 MRKKNLKMYKKAGEIKLKRKG FTLIEVVVVV AILGVIIAIA VPQVLKNINK         21
```

**Figure S1. Alignment of *Caldicellulosiruptor bescii* hypothetical proteins with putative pilin leader sequences.** The conserved glycine (G, purple) and glutamic acid (E, green) from the pre-pilin N-terminus cleavage domain (GFxxxE) are highlighted.

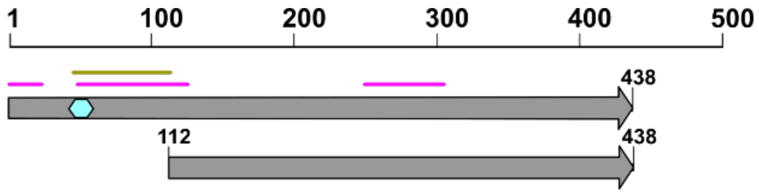


**Figure S2. Cloning strategy to produce soluble *C. bescii* PilA. A** truncation mutant of Athe_1880 was designed to remove the prepilin-type N-terminal cleavage/methylation domain-GFxxxE (cyan hexagon), as well as the highly hydrophobic region (magenta) and the predicted hydrophobic transmembrane domain (green). Ruler units are in base pairs.

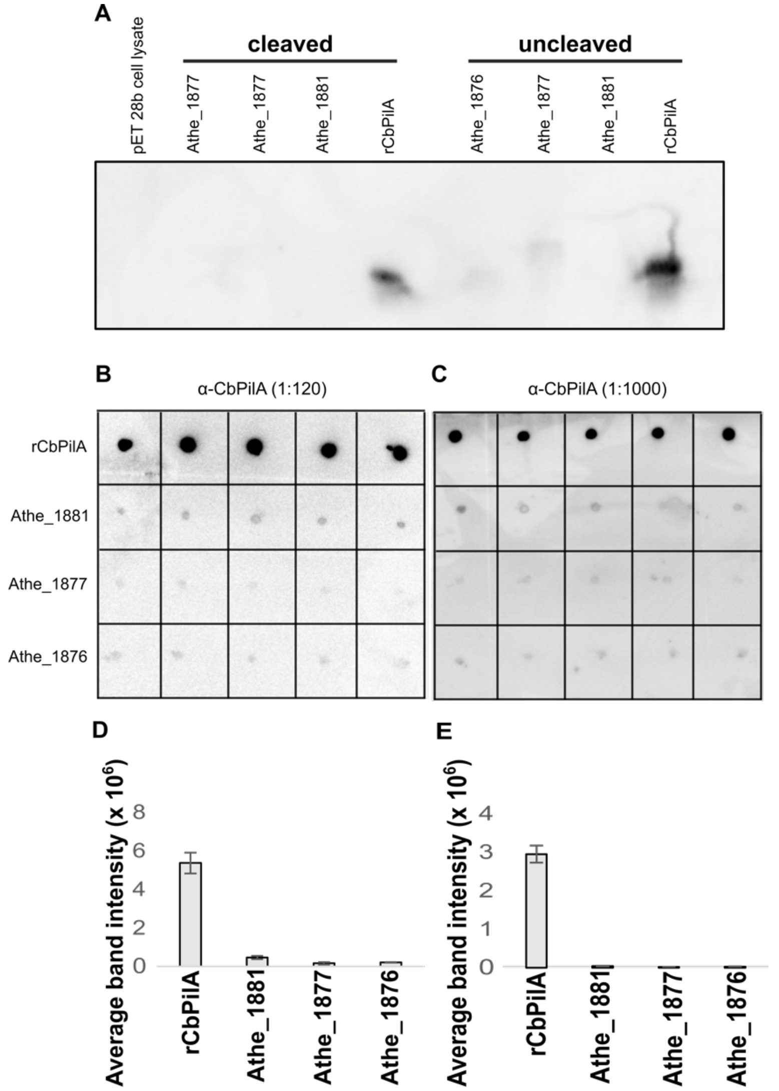


**Figure S3. Immuno-analysis of *C. bescii* pilins with anti-CbPilA.** (**A**) Immunoblot analysis of *C. bescii* pilins with anti-CbPilA. rCbPilA, Athe_1881, Athe_1877 and Athe_1876 were purified as described in the methods and the 6xHistags were cleaved using thrombin. Uncleaved as well as cleaved proteins were quantified, equalized to the same molar concentration (30 μM) and an equal volume of these pilins was loaded in each well. Heat treated *E. coli* cell lysate carrying an empty pET-28b expression vector was used as a negative control. (**B**) Immuno-dot blots probed using a 1:120 primary antibody dilution used for immunofluorescent microscopy. (**C**) Immuno-dot blots probed using an 1:1000 primary antibody dilution used for immuno-dot blots to quantify CbPilA expression. (**D**) Densitometry analysis from the immuno-dot blots in panel *B*. (**E**) Densitometry analysis for immuno-dot blots in panel *C*. For B and C, rCbPilA, Athe_1881, Athe_1877 and Athe_1876 were purified as described in the methods and 6xHis tags were cleaved using thrombin. Uncleaved as well as cleaved *C. bescii* pilins were quantified, equalized to the same molar concentration (30 μM) and an equal volume of these pilins was spotted on a PVDF membrane. For D and E, each column represents average intensity of the 5 replicates of each sample. Error bars represent standard error.

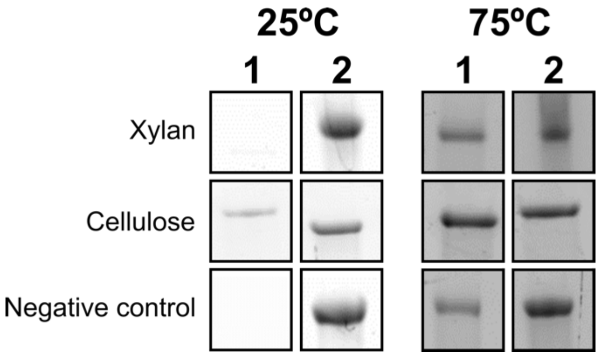


**Figure S4. Validation of the polysaccharide pulldown assay with recombinant *C. bescii* tāpirin.** (A) SDS-PAGE analysis of bound and unbound fractions of rCbtāpirin. **1**, bound fraction of rCbtāpirin; **2**, unbound fraction of rCbtāpirin. Polysaccharides tested were xylan and microcrystalline cellulose (Sigmacell). Negative control includes only rCbtāpirin without the polysaccharide. SDS-PAGE images are representatives of the results from assays run in triplicate. (B) Densitometry analysis for rCbtāpirin bound to cellulose at 75°C (C) Densitometry analysis for rCbtāpirin bound to xylan at 75°C. For B and C, each column represents average normalized intensity of the 3 replicates of each respective assay. Error bars represent standard error.

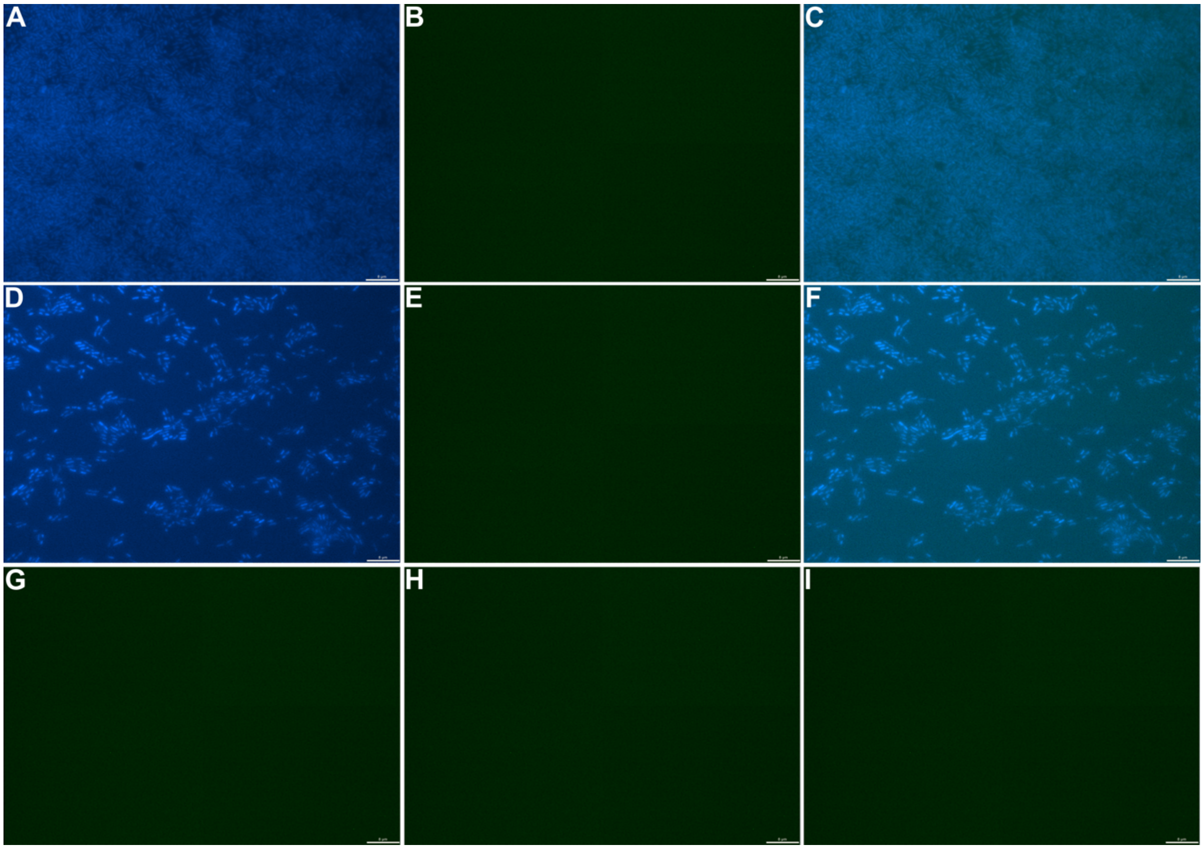


**Figure S5. Secondary antibody controls and biomass control for immunofluorescence detection of extracellular CbPilA from *C. bescii* cultures.** (A, B) DAPI, GFP and (C) color composite image of immunofluorescence microscopy of *C. bescii* cells grown on xylan and processed only with the secondary antibody. (D, E) DAPI, GFP and (F) color composite image of immunofluorescence microscopy of *C. bescii* cells grown on xylose and processed only with the secondary antibody. (G, H) DAPI, GFP and (I) color composite immunofluorescence microscopy of image of immunofluorescence microscopy of uninoculated LOD xylan medium processed with both primary and secondary antibodies. Brightness and offset of DAPI (blue) GFP (green) channels were balanced in the color composite images using Image-Pro Insight 9.1. Scale bars at the bottom of each image indicate 8μm.

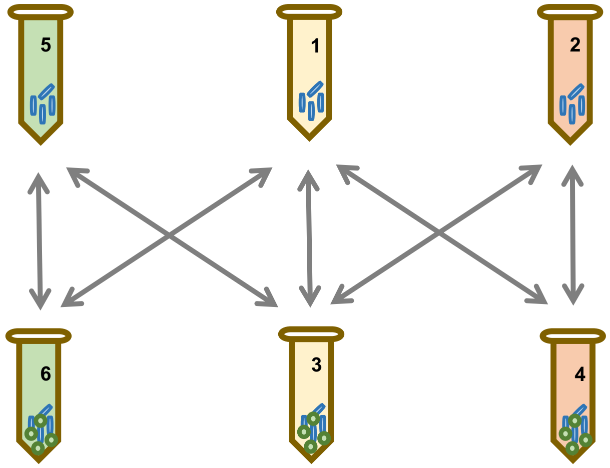


| | 1 | 2 | 3 | 4 | 5 | 6 |
|---|---|---|---|---|---|---|
| Cells | + | + | + | + | + | + |
| Polysaccharide | | | + | + | | + |
| rrCbPilA | | + | | + | | |
| BSA control | | | | | + | + |

**Figure S6. Experimental design for the *C. bescii* cell binding assay.** Blue rod: *C. bescii* cell, green circle: substrate. Four different treatment groups were set up to test the effect of two independent variables- substrate and protein. For the substrate variable the two treatments levels were: cells with binding buffer (50 mM sodium phosphate pH 7.2) alone and cells with binding buffer and substrate. For the protein variable the two treatment levels were, cells with rCbPilA alone and cells with rCbPilA and substrate.
